# Structure-Function Mapping of Olfactory Bulb Circuits with Synchrotron X-ray Nanotomography

**DOI:** 10.1101/2025.04.24.650439

**Authors:** Yuxin Zhang, Carles Bosch, Tobias Ackels, Alfred Laugros, Anne Bonnin, Jayde Livingstone, Carsten Waltenberg, Manuel Berning, Sina Tootoonian, Mihaly Kollo, Andrea Nathansen, Norman Rzepka, Peter Cloetens, Alexandra Pacureanu, Andreas T. Schaefer

## Abstract

Information is routed between brain areas via parallel streams. Neurons may share common inputs yet convey distinct information to different downstream targets. Here, we leverage the anatomical organisation of the mouse olfactory bulb (OB), where dozens of projection neurons (mitral and tufted cells, M/TCs) affiliate with a single input unit, a glomerulus^1–3^. To link functional properties of M/TCs to their anatomical glomerular association at scale, we combine *in vivo* two-photon (2P) imaging with synchrotron µCT^4–6^ anatomical analysis and targeted X-ray nano-holotomography (XNH)^7,8^. Improving XNH resolution for mm^3^ volumes enables us to reliably identify subcellular features, automatically segment >80,000 cell nuclei in individual experiments, and delineate several hundred functionally imaged projection neurons and their detailed morphology, including up to 20 M/TCs per individual glomerulus (“sister” cells). In over 2400 sister cell pairs, we consistently find that odour response amplitudes to a panel of 47 monomolecular odours are conserved between sister cells, with, however, distinct responses to individual odours. Responses correlated with anatomical features such as cell body position and lateral dendritic arborisation. Thus, sister cells neither simply relay glomerular inputs nor are they dominated by network activity. Instead, they show a “balanced diversity” in their responses, enabling efficient encoding of odour stimuli whilst retaining the overall structure of odour space. Thus, synchrotron X-ray tomography can reliably link subcellular anatomy to function in a non-destructive way across the mm^3^ scale. With recent advances in X-ray optics^9^ and the emergence of 4th generation synchrotrons^10,11^, it becomes conceivable to extend this highly accessible approach to entire brain regions with increasing resolution.

## Introduction

At its core, computation in the brain can be viewed as how neural circuits transform inputs and then output a new representation of information to downstream areas. Linking structure and function has long offered the promise to provide mechanistic insights into such computation^12–15^. In mammalian model systems, combining viral labelling of projections^16,17^ or synaptically connected neurons^18,19^ after *in vivo* imaging or *in vitro* electrophysiology^20,21^ has for example helped to elucidate structure-function relationships in sensory and motor systems. Combining *in vivo* imaging with *in vitro* electrophysiological recordings of connected neurons has provided evidence of how circuit connectivity relates to neuronal receptive field similarity^22–24^. As a recent alternative, all-optical-physiology approaches allow probing of physiology whilst measuring proxies of connectivity for a subset of neurons^25–27^.

Comprehensive anatomical analysis has so far often leveraged the resolution power of electron microscopy^28^. Advances in volume electron microscopy over the last decades have made it possible to densely delineate cellular composition and connectivity structure of neural tissues^28–35^. While pioneering work has combined these nano-connectomic approaches with prior functional imaging^36–40^, linking these two modalities is often challenged by the relevant scales involved. Electrons penetrate tissue only for a few 100s of nanometers^41,42^, necessitating tissue sectioning or ablation for the acquisition of volume information^31–33,43–45^. This makes it challenging to routinely access volumes comparable to mammalian brain regions that span the mm^3^ range. Thus, pioneering volume EM studies combining structural with prior functional analysis are still rare and - with very few exceptions^40^ - have been restricted to the analysis of an individual sample and volumes of typically less than 0.1 mm^3^. X-ray tomography has emerged as a promising and complementary addition to the toolbox for volume anatomical analysis of brain tissue^4–6,8^. It leverages the fact that X-rays readily penetrate tissues over mm or cm. Illuminating samples with X-rays of wavelengths typically between 0.01 nm and 1 nm (∼1keV-100keV) allows recording of projections with the transmitted light, using X-ray absorption or scattering to generate contrast^46^. Hundreds to thousands of projections are recorded at different rotation angles and their combination results in a 3-dimensional tomogram, obtained without any physical sectioning or ablation. Contemporary synchrotrons provide highly coherent and intense X-ray beams^41,42,47^. Synchrotron X-ray computed tomography with propagation-based phase contrast (SXRT) can be used to obtain structural information for volumes of several mm^3^ to cm^3^ and can provide 3d histology resolving anatomical layers, cell bodies and large neurites^4–6^. Other phase contrast techniques such as X-ray ptychography^48–50^ or holography^7^ can provide substantially increased resolution (as little as <5 nm in non-biological samples^51^). For neuronal tissue, X-ray nano-holotomography (XNH, <100 nm resolution^8^) or ptychography(<40 nm resolution^9^) have shown promise to resolve subcellular features. XNH in particular is a full-field technique where X-ray optics are used to focus the beam^52–54^, recording a geometrically magnified image of the specimen. By combining projections from different propagation distances, a phase map can be reconstructed^7,55^, allowing for rapid acquisition of subcellular-resolution data.

The mouse olfactory bulb (OB) provides a powerful model system to leverage the capabilities of X-ray tomography to delineate structure and relate it to functional measurements to investigate how neural circuits transform sensory representations. Olfactory receptor neurons (ORNs) in the nose detect smells and send their axons into the OB where ORNs that express a common receptor converge onto a glomerulus, a distinct ∼100 µm spherical neuropil structure^56^. There, projection neurons (PNs) receive ORN inputs with their ∼µm thick apical dendrites and relay information, shaped by numerous local interneurons, to a set of downstream areas^1,2,57^. Each glomerulus acts as a functional module and is associated with a dedicated set of ∼20-50 PNs, separable into deeper mitral cells (MCs), projecting to e.g. piriform cortex, cortical amygdala, entorhinal cortex, olfactory tubercle and more superficial tufted cells (TCs), exclusively projecting to rostral brain areas^1,2,57^. Sharing overlapping projection targets, mitral and tufted cells (M/TCs) also display similar physiology, yet MCs are viewed to be more shaped by local circuitry^2,58–60^. How the dozens of “sister” M/TCs associated with an individual parent glomerulus differ, how they might each represent a differently processed version of the same excitatory input, remains unclear, largely because PNs belonging to a specific glomerulus are distributed across 100s of micrometers to millimetres ^3,61^. Optogenetic approaches have been pioneered to identify individual pairs of putative sister cells^62,63^ associated with one specific genetically labelled glomerulus^64^ as have targeted electrical recordings from an individual exogenous glomerulus^65^, or electroporation approaches^61^, suggesting a complex relationship within sister cell activity.

The molecularly tight organization of the olfactory bulb input layer^66^ has often led to the notion that M/TCs relay information about the molecular identity of the parent glomerulus and its associated receptor. In its extreme, this predicts stereotypical sister M/TC responses to the entire panel of odours activating the parent glomerulus thereby maintaining the molecular receptive range of the parent glomerulus, relaying chemical similarity faithfully to downstream areas **(Supp Fig 5_7a1-e1)**. On the other hand, the OB network is dominated by local computation with strikingly 90% of OB neurons being local interneurons^67^. This in turn suggests that M/TC activity is overwhelmingly dominated by network effects predicting high diversity of sister M/TCs and concomitantly high coding capacity with minimal redundancy **(Supp Fig 5_7a3-e3)**. Addressing these predictions necessitates recording physiological responses and unambiguously identifying sister M/TCs at scale across individual animals.

To address this outstanding question, here, we extend X-ray tomography imaging to the scale necessary for this circuit analysis. We show improvements in XNH acquisition speed and signal-to-noise as well as new image reconstruction techniques to extend the accessible volumes with subcellular resolution to the mm^3^ scale. We leverage these developments to combine targeted XNH anatomical analysis with large-volume SXRT and prior functional imaging *in vivo* at scale. This makes it possible to anatomically identify several hundred functionally analysed M/TCs in a single experiment with their glomerular affiliation, resulting in more than a thousand pairs of anatomically identified sister cells in a single specimen. Moreover, the robustness of the approach made it possible to collect three biological replicates. Consistently, we find that the population of sister cells preserves odour identity information relayed from a glomerular module. Sister cells differ, however, in responses to specific odours and in their dynamics, shaped by somatic location and dendritic anatomy.

Combining in vivo imaging with XNH anatomy at scale, we provide a comprehensive description of how inputs to a brain region are transformed and broadcast downstream. We find that the OB network implements a finely balanced diversity, allowing for efficient encoding of odours whilst maintaining the similarity structure of odour space **(Supp Fig 5_7a2-e2)**.

## Results

### Linking function to structure with XNH

To assess the functional properties of sister cells and, more broadly, to link structure to function, we first performed 2P imaging experiments *in vivo*, genetically expressing GCaMP6f in M/TCs and presenting a panel of 47 odours **(Figure 1a, Supp Fig 1_1,** both M/TC somata and glomerular signals were imaged, see **Supp Fig 1_9)**. To assess the base state of the early olfactory system in the absence of specific, e.g. attentional contexts, we performed these experiments in anaesthetised mice, and recorded all experiments in an equivalent region of the OB around a fluorescently-labelled genetically-identified glomerulus (MOR174/9)^1,3^. After functional imaging, we extracted the OB, sectioned it and stained it with a high-contrast metal staining protocol (rOTO, see Methods). After embedding, we assessed overall structural integrity using a benchtop µCT scanner **(Supp Fig 1_8)**. Subsequent SXRT gave detailed anatomical delineation of glomeruli, layers and cell bodies **(Supp Fig 1_2)**. This allowed for further quality control and for targeting the very region functionally imaged *in vivo* using the blood vessel pattern and other histological features conserved in both datasets, such as borders of glomeruli, as landmarks. To optimise quality for XNH, we then extracted a ∼1 mm diameter x 600 μm height cylindrical pillar using a recently developed fsLaser milling approach^68^ **(Supp Fig 1_3)**. XNH is a cone beam technique providing substantial geometric magnification and resolution down to <100 nm as pioneered in beamline ID16A at the European Synchrotron **(Supp Fig 1_4)**. Whilst a (rapid) full-field technique, in its current implementation at ID16A, the field of view (FOV) is restricted to ∼200 µm. Thus, to acquire mm^3^ volume, we adopted a tiling approach, acquiring a set of up to 36 partially overlapping tiles covering the entire relevant tissue volume **(Figure 1b** and **Supp Fig 1_4)**. We employed continuous scanning to minimise artefacts associated with interior tomography and speed up data acquisition^69^. To further avoid ring artefacts and reduce structured noise, we developed an approach that used two independent, spatially offset projections computationally combined using a machine learning protocol **(Supp Fig 1_4**, see Methods and ref. ^70,71^). With all these improvements and the recently upgraded Extremely Brilliant Source at ESRF, acquiring an individual tile required ∼110 minutes, resulting in 66 hours of total imaging (and motor movement) time for all 36 tiles. Individual (virtual) tiles were then computationally stitched together to yield a continuous, artefact-free mm^3^ volume where glomeruli and nuclei could be segmented automatically **(Figure 1b, 2c2** and **Supp Fig 1_6**, ref. ^72^). The ease of sample preparation without sectioning and the fast speed of acquisition allowed us to acquire samples from three individual animals.

**Figure 1:**
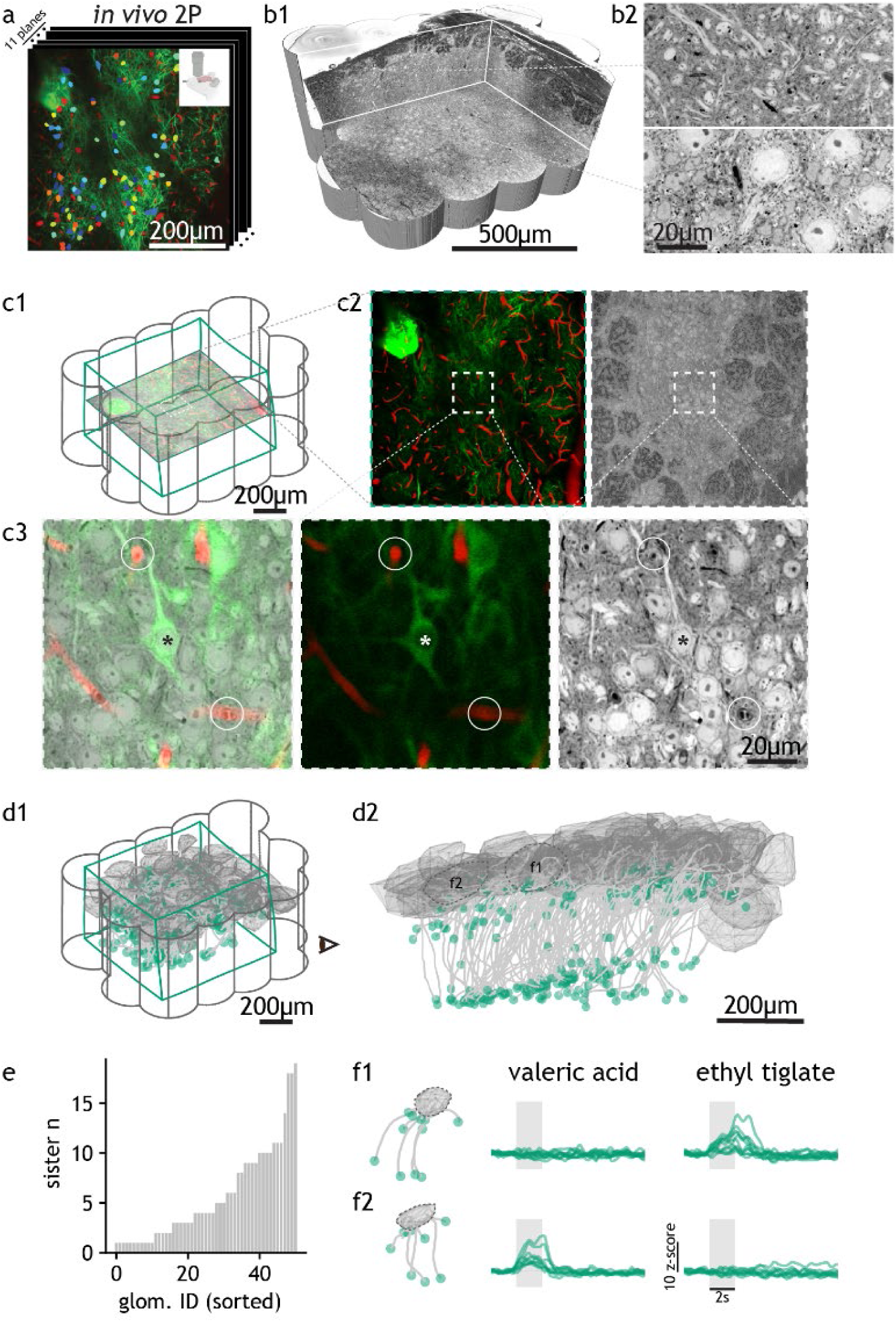
Correlative *in vivo* functional imaging and X-ray nano-holotomography (XNH). **(a)** Example plane from *in vivo* 2P Ca^2+^ imaging of mouse olfactory bulb with coloured masks overlaid on functionally imaged cells (tufted cell plane). **(b)** XNH dataset volume **(b1)** with example images of external plexiform layer dendrites (top) and mitral cell somata (bottom) **(b2)**. **(c)** Precision of 2P-XNH dataset correlation. **(c1)** XNH (grey) and 2P (green) datasets cover the same volume of tissue. **(c2)** Example 2P plane (green: GCaMP signal, red: blood vessel signal) and its corresponding XNH plane. **(c3)** Zoomed-in field of view showing precisely overlaid cells (asterisks) and blood vessels (circles) in the combined 2P-XNH and individual 2P and XNH datasets. **(d)** Apical dendrite tracings of functionally imaged projection neurons (green dots, n = 289) to their parent glomeruli (grey hulls, n = 50) viewed *in situ* **(d1)** and from one side **(d2**). **(e)** Counts of sister cells per glomerulus with glomeruli sorted by increasing number of associated sister cells. **(f)** Sister cells from two example glomeruli and their responses to two odours, with the odour presentation period shaded in grey (sister n = 11, 9). See also **Supp Video 3** and https://github.com/yz22015/sister_2p_xnh/tree/main.

### Odour tuning of sister cells at scale

To align *in vivo* imaging to anatomy, we first used general anatomical landmarks such as glomeruli. We created fine-grained additional landmarks by labelling vasculature *in vivo*^73^ for more detailed alignment. Iterative alignment resulted in a precision of a few µm, reliably allowing identification and precise localisation of individual neurons imaged in vivo **(Figure 1c,d)**. To probe odour representation across sister cells, in the *in vivo* imaging experiments, we presented a panel of 47 odours, broadly spanning chemical space **(Supp Fig 1_7, Supp Table 1)**. As expected, this panel of odours elicited broad and diverse responses across the population of projection neurons **(Figure 2a1)**. To assess how glomerular affiliation might inform these response patterns, we sought to identify M/TCs in the corresponding XNH volume and trace their apical dendrites **(Supp Fig 1_4, Supp Video 1, 2)** to the respective parent glomeruli. We identified 289 M/TCs (162 TCs, 127 MCs) associated with 50 glomeruli with up to 20 sister cells for a single glomerulus **(Figure 1d,e)**. Seven glomeruli had more than 10, and 20 glomeruli had more than 5 sister cells each with verified anatomy and detailed functional profiles **(Figure 1e)**. This allowed us to sort the odour response patterns of MCs and TCs by their parent glomerulus. For example, in one glomerulus (glomerulus 3), all 11 sister TCs and MCs robustly responded to ethyl tiglate while not responding to valeric acid (activity integrated over a 2-second window). In a second glomerulus (glomerulus 33), all 9 simultaneously recorded sister cells responded robustly to valeric acid but not to ethyl tiglate **(Figure 1f)**.

**Figure 2:**
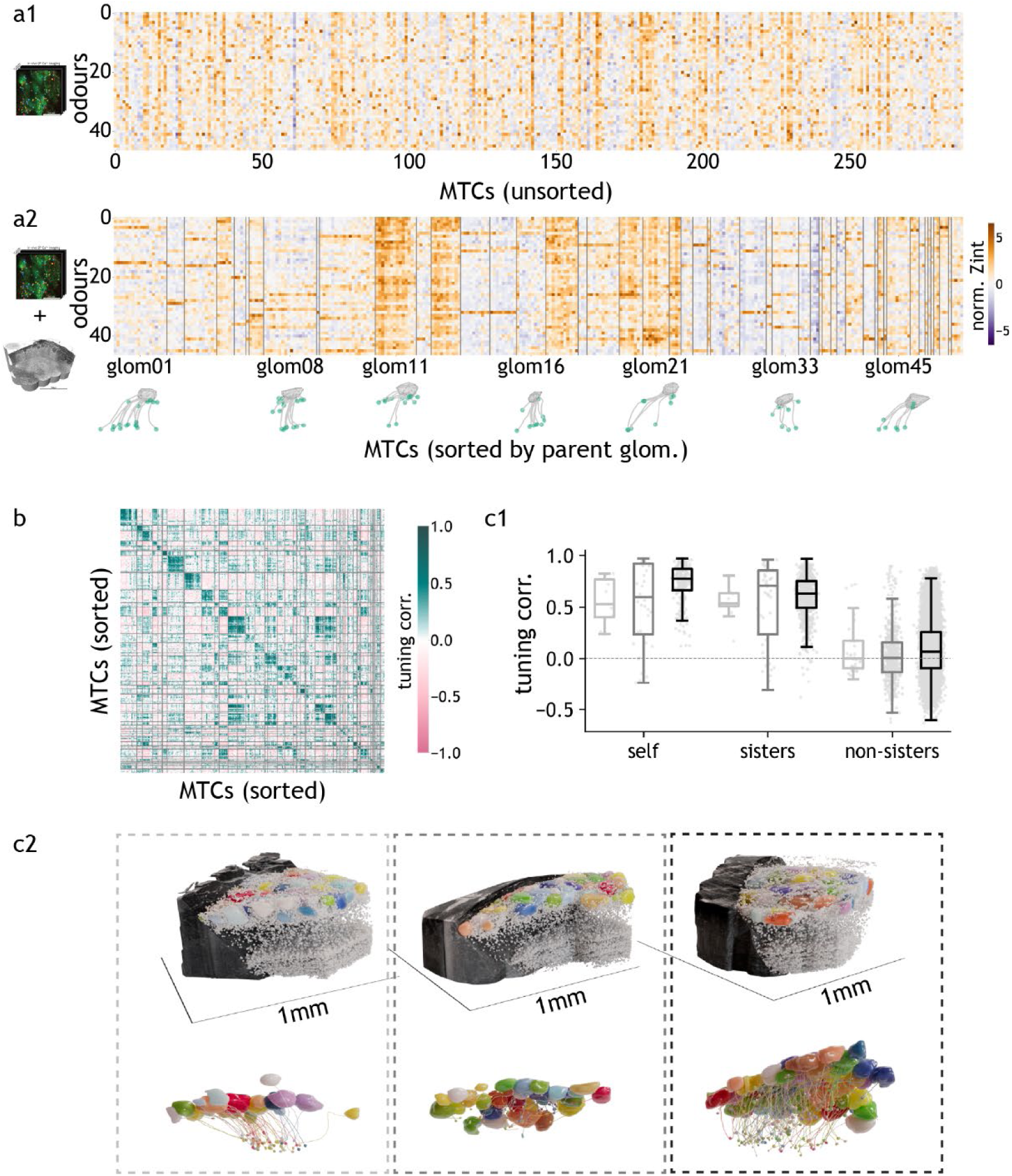
Odour tuning similarities of anatomically identified sister cells. **(a)** Heatmap of odour response integral of cells, without sorting, equivalent to having the 2P dataset alone **(a1)** and with cells sorted by parent glomerulus identity, enabled by apical dendrite tracing in the XNH dataset **(a2)**. Response integral (‘Zint’) defined as dF/F z-score integral over the odour presentation period (2 s), averaged across all repeats (n = 4) and normalised by the standard deviation of responses to all odours per cell. **(b)** Heatmap of split-repeat odour tuning correlation of cells, sorted by parent glomerulus identity (see Methods). **(c)** Tuning correlation summary across three datasets. **(c1)** Boxplot of tuning correlation values (7 s integral) for each cell across repeats (‘self’) and for sister and non-sister cell pairs, for three datasets independently acquired from three animals (box = first and third quartile, midline = median, whiskers = most extreme, non-outlier data points. For the light grey, dark grey and black samples, self, sister and non-sister mean values are [0.558, 0.477, 0.073], [0.535, 0.486, 0.03] and [0.728, 0.613, 0.086], respectively and sister > non- sister one-tail t-test *p* values are 1.37E-07, 4.34E-11 and 6.49E-146, respectively). **(c2)** The corresponding XNH datasets with segmented glomeruli and nuclei *in situ* (top) and distilled apical dendrite tracing (bottom). All functionally imaged cells that were traced to their parent glomeruli were used in the analyses of this figure. For **c1**, only glomeruli with > 1 sister cell and only strictly responding cells were included (see Methods). See also **Supp Videos 3, 4.**

Across the population of 289 simultaneously recorded MCs and TCs, no clear structure was apparent in their responses to the panel of 47 odours **(Figure 2a1)**. Sorting by anatomical affiliation, however, revealed high consistency in the (time-averaged) responses of sister cells: Cells belonging to the same parent glomerulus showed highly correlated response profiles (response averaged over 2 seconds, **Figure 2a2, b, Supp Fig 2_1**). Cells responded largely stereotypically, with responses almost as similar as between repeated trials (r=0.52 for repeated trials vs r=0.40 for sister cells **Figure 2c1**). Non-sister cells, on the other hand, showed no correlation in responses on average (r=0.012, sister > non-sister 1-tail t-test p = 2.43e-115, **Figure 2c1**, see **Supp Fig 2_2** for rare examples of non-sister cells sharing high tuning correlation) and tuning correlation between different glomeruli was weak and largely independent of their relative position **(Supp Fig 2_3)**. This significant stereotypy in sister cell tuning was highly consistent between all three biological replicates (**Figure 2c1,c2, Supp Fig 2_1**). Thus, under baseline conditions in anaesthetized mice, network computation does not dominate the transformation between input and output in the olfactory bulb. This consolidates what was suggested by pioneering studies based on recordings from individual pairs of sister cells or individual glomeruli ^61,62^. Instead, the output (MC and TC responses) retains key response features of the input (approximated here by the average activity of the parent glomerulus, **Supp Fig 3_2d-f**).

### Common sister cell responses to strong and weak odours

What defines the high tuning correlation of sister cells? Do sister cells only share similar response profiles to a few common odours, or are response patterns consistently mirrored across the entire panel of 47 odours? To test this, we first probed responses only to a small set of odours that engaged the parent glomerulus most strongly. Sorting sister cell odour responses by the parent glomerulus showed a consistent pattern **(Figure 3a)**: Those odours that engaged the parent glomerulus most strongly (see Methods) consistently also engaged the corresponding sister cells. In fact, the overlap between the identities of the most potent odours was nearly as high between sisters as between repetitions for an individual cell **(Figure 3b1,b2)**. Only when considering 10 or more odours did sister cell overlap become notably weaker than overlap for repetitions **(Figure 3b2)**. Does that imply that sister cells do not share response patterns to “weak” odours? We removed the responses to the 5 strongest odours from the dataset to assess this and calculated the tuning correlation between cell responses and their parent glomerulus. Indeed, tuning correlation was reduced compared to the whole odour set (**Figure 3c**, compare “all” to “weak”). This difference, however, was slight and cells were vastly more strongly correlated to the parent glomerulus than to other glomeruli (**Figure 3c**, compare “weak” to “other”). Even when removing the 10+ strongest odours, sister cell responses correlated highly with their parent glomerulus **(Figure 3d)**. Similarly, removing the strongest odours and calculating cell-cell tuning correlation showed that sister cells robustly share tuning to weaker odours as well **(Supp Fig 3_1)**. Thus, sister cells relay information about the parent glomerular response to downstream targets for both strongly and weakly activating odours.

**Figure 3:**
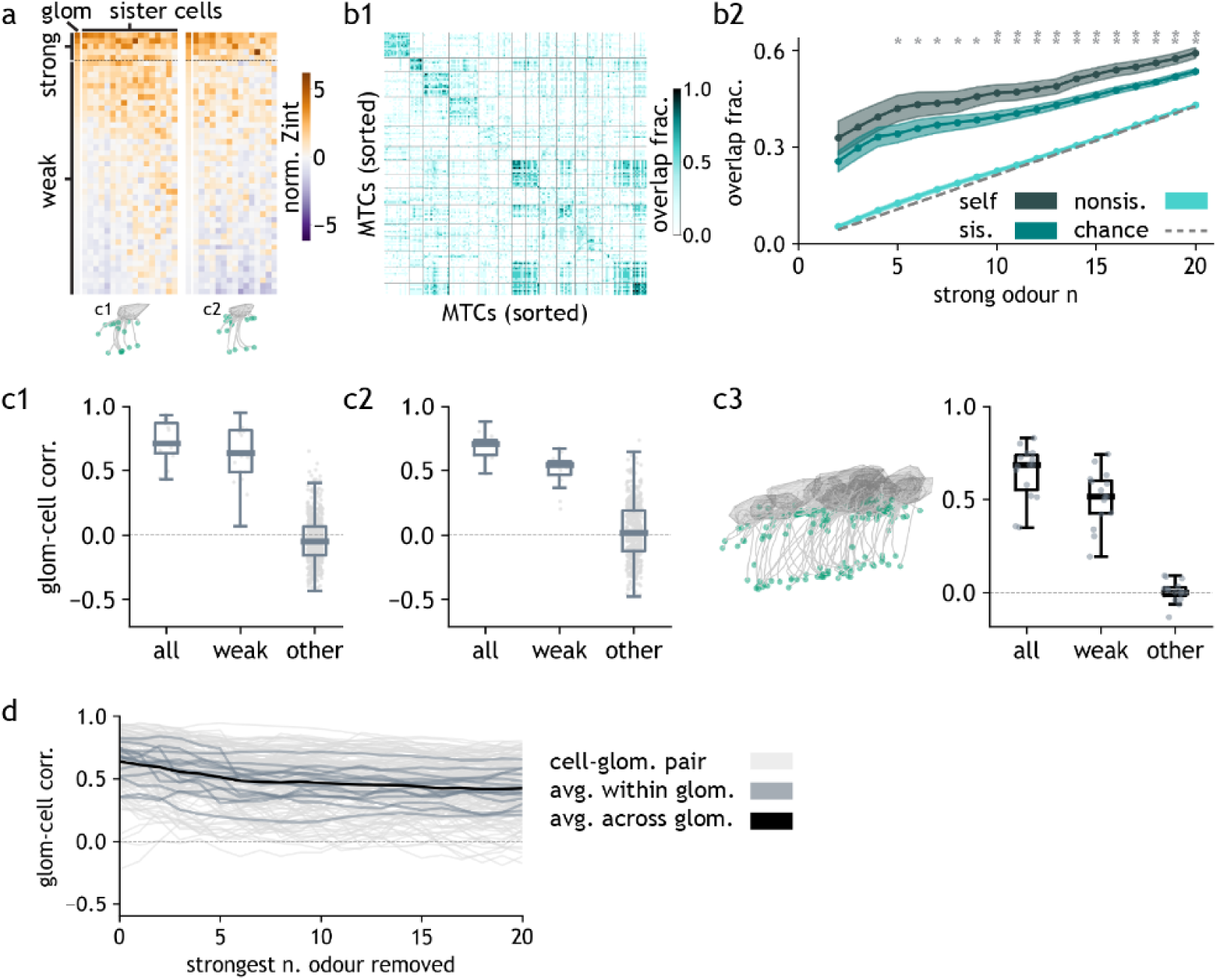
Sister cell responses to strongly and weakly activating odours. **(a)** Heatmap of odour response integral of sister cells from two example glomeruli. Odours ranked by the response strength of the parent glomerulus. Unless specified, top n = 5 odours considered strong and the rest weak by convention. **(b)** Similarity in strong-odour identity across cells. **(b1)** Heatmap of strong-odour overlap among cell pairs (see Methods), with cells sorted by parent glomerulus identity. **(b2)** The mean and 95% confidence interval of strong-odour overlap for individual cells across repeats (‘self’) and for sister and non-sister cell pairs, as a function of increasing number of strong odours considered. Dashed line represents chance-level overlap averaged across 1000 simulations of randomly drawn odours. ‘sis’ > ‘nonsis’ one-tail t-test *p* < 0.001 (*) or < 10^-5^ (**). **(c)** Boxplot of tuning correlation between glomeruli and cells, for two example glomeruli **(c1-2)** and mean values pooled across all glomeruli **(c3)**. Three categories represent tuning correlation between parent glomeruli and its sister cells, calculated using all odours (‘all’) or only weak odours (‘weak’); or between random non-parent glomeruli and cells using weak odours (‘other’). Box = first and third quartile, midline = median, whiskers = most extreme, non-outlier data points. For **c1-3**, the mean values for ‘all’, ‘weak’ and ‘other’ categories are [0.71, 0.628, -0.035], [0.683, 0.497, 0.033] and [0.673, 0.507, 0.004], respectively and the ‘weak’ > ‘other’ one-tail t-test *p* values are 5.29E-45, 1.21E-15 and 5.61E-13, respectively. **(d)** Glomerulus-cell tuning correlation after removing n strong odours, for all parent glomerulus-sister cell pairs, the mean per glomerulus and the mean across all glomeruli. Only glomeruli with 2P data and > 5 sister cells and only strictly responding cells (see Methods) were used in the analyses of this figure.

### Diversity of sister cell responses

Does this imply that sister cells simply relay redundant information to downstream targets? Whilst overall tuning correlations are indeed high, responses can differ significantly for individual odours and cells. For example, sister cells can share identical responses to one odour **(Figure 4a top row)** whilst showing opposite polarity **(Figure 4a1)** or different time course **(Figure 4a2,a3)** to another odour (see also **Supp Fig 4_1**). To quantify these distinct differences, we calculated the similarity of the sister cell responses for every glomerulus and odour (see Methods). For about a third of odour-glomerulus pairs, the time course of sister cell responses was highly similar (correlation > 0.9, “stereotypic”). For another third, responses were diverse (correlation <0.7), with the remainder showing intermediate similarity **(Figure 4b)**. Approximately half of the glomeruli displayed stereotypical responses to odours **(Figure 4c1)**. This was largely independent of glomerulus size, but the larger the number of sister cells recorded from a glomerulus, the smaller the fraction of stereotypical odours (**Supp Fig 4_1c**, r=-0.594, p=0.001). Conversely, most odours evoked stereotypical responses in at least some glomeruli **(Figure 4c2),** and odour response stereotypy was independent of the chemical class of the odour **(Supp Fig 4_1d)**. What does this suggest for cell and glomerular identity? A simple linear classifier could not only predict parent glomerular identity (**Figure 4e2**, 74.1%, 3.8% chance), a classifier trained on individual cells performed substantially above chance, identifying the correct cell in 32% of cases (**Figure 4d,e1**, 0.4% chance). Consistent with the high tuning correlation between sister cells, misclassifications typically happened within a glomerulus for sister cells **(Figure 4d2)**. Consequently, odour encoding was more accurate with an increasing number of sister cells **(Supp Fig 4_2c)**. Thus, we have established that sister cells do not simply relay glomerular information to downstream regions but that individual sister cells reproducibly differ in distinct odour response patterns (in time course, strength or even polarity, in agreement with earlier work investigating individual sister cell pairs^62^), thereby increasing encoding prowess.

**Figure 4:**
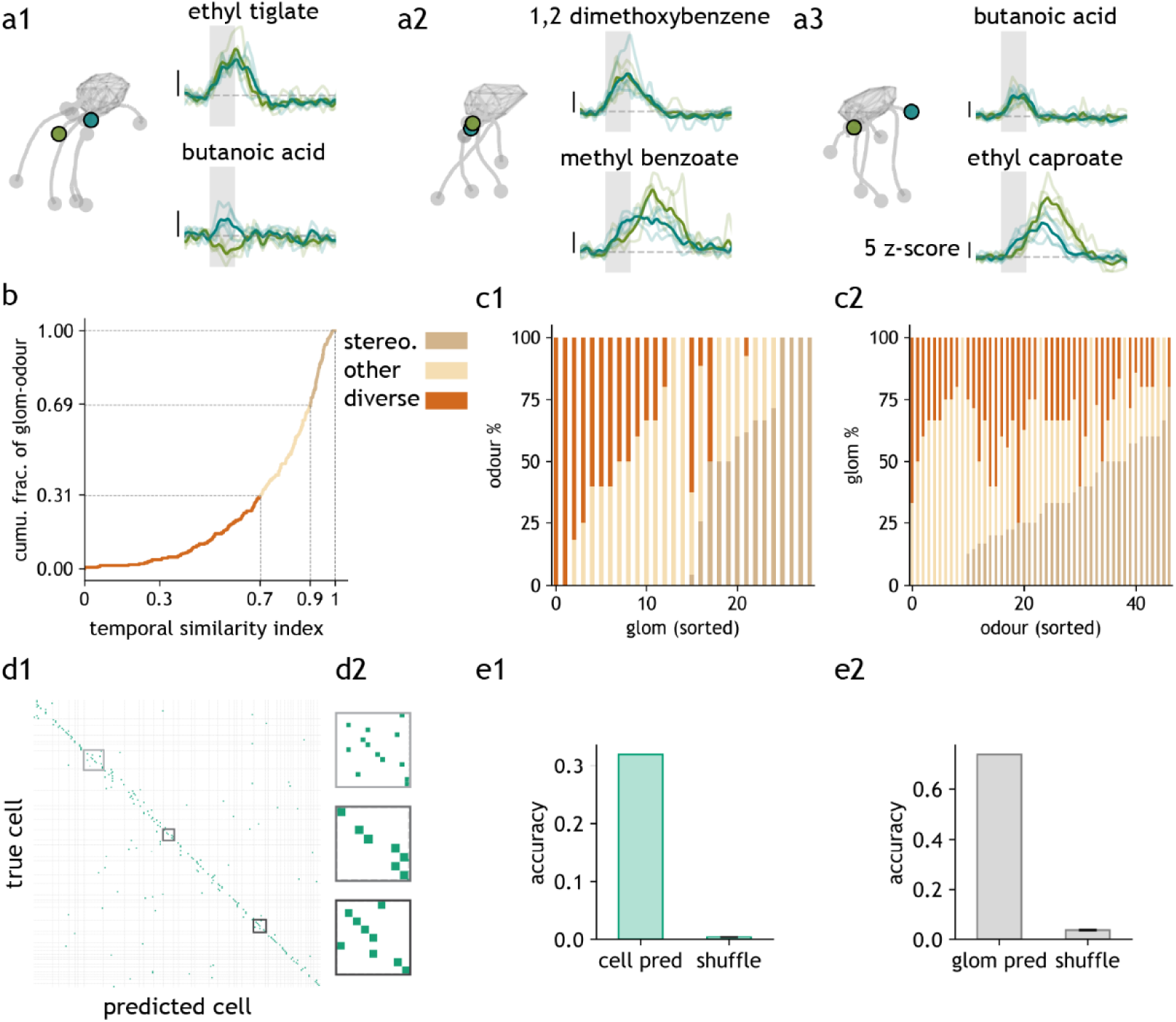
**Temporal differences in sister cell responses.** **(a)** Comparison of sister cell responses to different odours. Raw dF/F z-score traces averaged across repeats (dark lines) and for each individual repeat (light lines) of a sister cell pair (shades of green), responding to odours that elicited stereotypical response (top) or diverse response (bottom), for three example glomeruli **(a1-3)**. **(b)** Cumulative histogram of temporal similarity index (see Methods) of all glomerulus-odour pairs. An index < 0.7 is considered diverse and ≥0.9 considered stereotypical. **(c)** Fraction of stereotypical or diverse glomerulus-odour pairs, summarized per glomerulus **(c1)** or per odour **(c2)**. **(d)** Cell prediction using a linear SVM (see Methods). **(d1)** Confusion matrix of prediction result, with cells sorted by parent glomerulus identity. **(d2)** Zoom-in view of three example glomeruli from **(d1)**. **(e)** Bar plots of prediction accuracy for correctly identified cells (0.32) **(e1)** or parent glomeruli (0.74) **(e2)** compared to chance level (shuffled training labels, 0.004 for cells and 0.038 for glomeruli). Only glomeruli with multiple sister cells and only strictly responding cells (see Methods) were used in the analyses of this figure. For **(b-c)**, only odours that reliably elicited responses in at least 1 sister cell per glomerulus were included.

### Anatomical correlates of diversity

What enables this diversity in responses? So far, we have used the anatomical information provided by the XNH imaging to identify apical dendrites and, thus, parent glomeruli reliably and at scale. High-resolution XNH imaging, however, also allows us to identify e.g. soma position and its relation to the parent glomerulus. Indeed, comparing somatic odour responses to odour activation in the parent glomeruli reveals that the further a soma is from its parent glomerulus, the less similar responses are (**Figure 5a1,b1**, r = -0.385, p = 1.51x10^-8^**, Supp Fig 5_3** see also ref ^61^). Projection neurons in the OB fall into several classes; in the first instance, into mitral (MC) and tufted (TC) cells. MC somata are situated in a narrow band deeper in the OB, whereas TC cell bodies are located more superficially. The trans-genic line we employed (Tbet-Cre) is thought to label tufted and mitral cells without any known further specificity. Separating along the TC/MC divide shows that this correlation persists within the TC (**Figure 5a2, b2**, r = -0.173, p = 0.0418) and, in particular, MC (**Figure 5a3, b3**, r = -0.258, p = 0.0412) population (see also ref ^61^). Importantly, as a whole, the TC population was more similar to the parent glomerulus activity **(Figure 5c,d)** and sister cell correlations between TCs were higher than between MCs **(Supp Fig 5_1a-d,** for TC-MC comparison, see **Supp Fig 5_1f)**.

**Figure 5:**
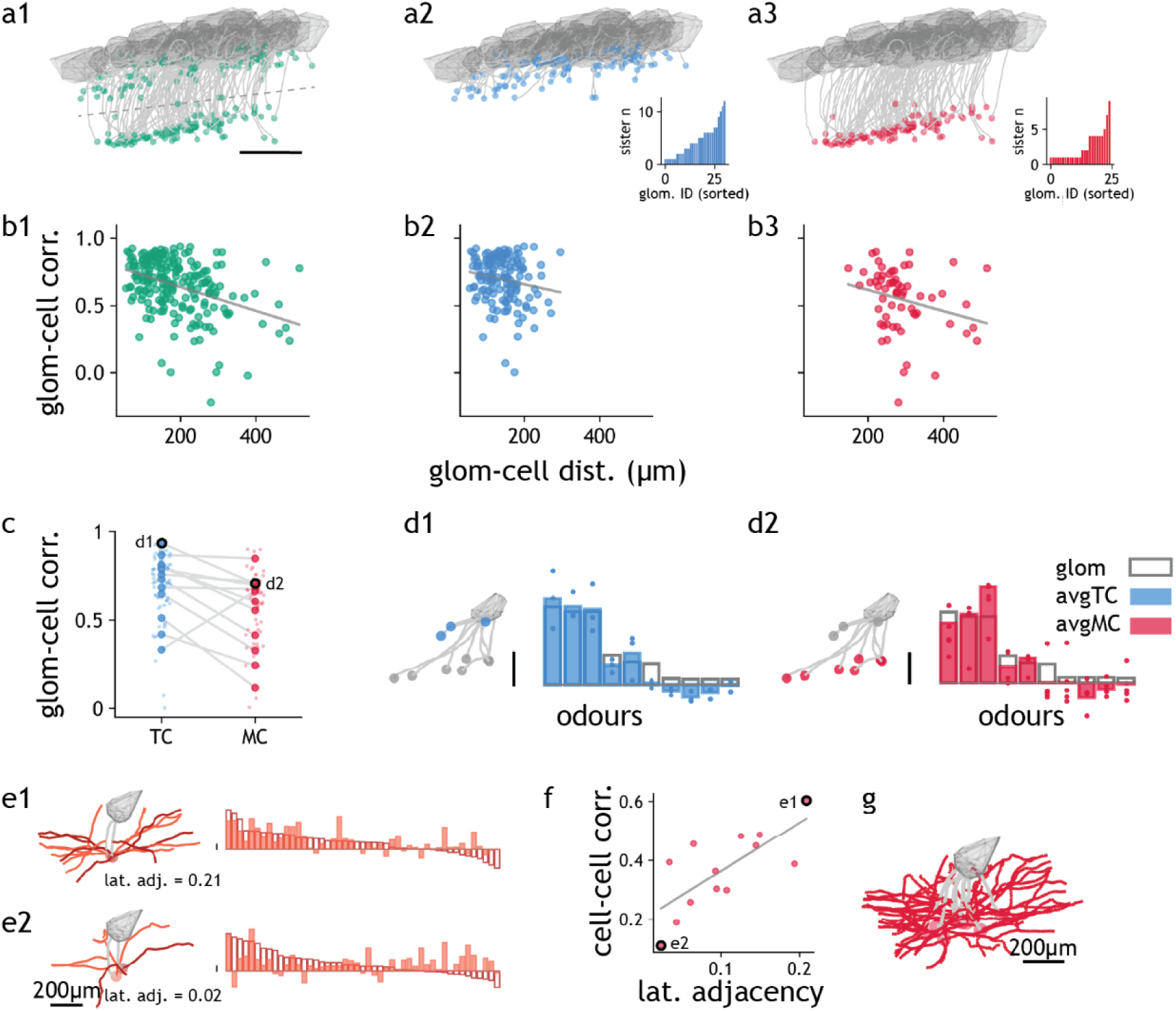
Anatomical correlates of sister cell tuning. **(a)** Division of projection neurons into TCs and MCs by soma depth **(a1)**. Apical dendrite tracing and count histogram of TCs **(a2)** and MCs **(a3)** per glomerulus. **(b)** Parent glomerulus-cell odour tuning correlation against glomerulus centroid-soma centroid distance, for all projection neurons **(b1)**, TCs **(b2)** and MCs **(b3)**. For **b1-3**, correlation coefficient values are -0.385, -0.173 and -0.258, respectively and *p* values are 1.51E-08, 0.042 and 0.041, respectively. **(c)** Comparison of tuning correlation between parent glomerulus and TC (blue) or MC (red), with small dots representing individual glomerulus-cell pairs and large dots representing the average across all sister cells per glomerulus. **(d)** Example odour response integral of sister TCs **(d1)** and MCs **(d2)**, with dots representing individual cells and coloured bars representing the average within cell types, overlaid with that of the parent glomerulus (grey bar). **(e)** Apical (grey) and lateral (coloured) dendrite tracings of example MC pairs with high **(e1)** and low **(e2)** lateral adjacency score and their response integral bar plot. **(f)** Relationship between odour tuning correlation against lateral adjacency score for MC pairs from an example glomerulus (*r* = 0.726, *p* = 0.005). **(g)** Apical and lateral dendrite tracings of MCs of the example glomerulus in **(e-f)**. **(a)** only includes glomeruli with 2P data and their strictly responding cells (see Methods, n = 139 TCs, 63 MCs). **(b, c)** contain those glomeruli with > 1 TC and > 1 MC.

How does the network shape sister cell activity? While a significant amount of computation is thought to occur within the glomerular circuitry^58,74–78^, granule cells have long been thought to implement various key computations shaping the output of the OB. Granule cells form dendrodendritic, often recurrent, synapses with the lateral dendrites of MCs and TCs. Thus, we utilised the XNH dataset again to reconstruct the full lateral dendritic tree of MCs and TCs **(Figure 5e-i, Supp Fig 5_4)**. To assess the potential for GCs to modulate activity in pairs of sister cells, we calculated the physical proximity of lateral dendrites from pairs of cells (reasoning that for two MCs to contact the same set of GCs and thus share inhibitory modulation, they would need to sample overlapping space, see Methods). Indeed, such a “lateral adjacency score” in glomerulus 10 predicted the similarity in odour responses: MCs with closely adjacent lateral dendrites had more similar odour responses **(Figure 5e-g)**. While this is consistent with a key role in lateral dendrites (and likely GCs) in shaping odour responses, the strength of this link varied between glomeruli **(Supp Fig 5_2)**, indicating that a multitude of mechanisms might underlie the diversity of sister cell responses.

## Discussion

Here, we have leveraged synchrotron-based X-ray nano-holotomography (XNH) alongside *in vivo* two-photon (2P) functional imaging to map the anatomical and functional properties of mitral (MC) and tufted (TC) cells in the mouse olfactory bulb (OB) at an unprecedented scale. This approach enabled us to simultaneously record from several hundred anatomically verified projection neurons, including up to 20 neurons from individual glomeruli, yielding a dataset of over 2,400 sister cell pairs across three biological replicates. Our findings show that while odour tuning among sister cells is strikingly conserved, individual responses display heterogeneity in response dynamics. Furthermore, MCs exhibit greater diversity in their response properties compared to TCs, a difference that can be partially explained by their anatomical distribution and network interactions. Together, this suggests a model of “balanced diversity” where the output of the OB efficiently encodes odour space whilst maintaining chemical similarity structure **(Supp Fig 5_7)**.

### Odour Coding in Sister Cells: Between Redundancy and Diversity

Our findings provide a comprehensive perspective on models of olfactory processing. Sister cells do not merely act as redundant relays, nor are they completely independent in their representations. Instead, our data suggest a balance between preserving odour identity and maximising information encoding (**Supp Fig 5_7)**. The strong conservation of tuning across sister cells ensures that odour identity information is robustly transmitted to downstream targets, while their heterogeneity in response dynamics increases the encoding capacity of the OB compared to a pure relay model.

Mechanistically, this conserved coding likely arises from the strong feedforward drive imposed by olfactory receptor neuron (ORN) input, ensuring that odour identity remains stable across multiple downstream projections. At the same time, heterogeneity within the sister cell population, likely a result of integrating information across the OB, enables differential contributions to sensory processing. This in turn ensures that downstream regions receive both a faithful representation of odour identity and similarity structure as well as additional dimensions of information.

### Mitral vs. Tufted Cells: Functional Differences in Odour Processing

Our data reveal a systematic difference between MCs and TCs in their odour representations. MCs display greater decorrelation in their responses compared to TCs. This aligns with the hypothesis that TCs provide a more immediate, feedforward representation of odour information, while MCs—through deeper integration into the OB’s inhibitory network— contribute to a more processed, context-dependent output^58,59^. This specialisation is likely critical for enabling distinct computations across olfactory recipient regions: TCs predominantly target rostral brain areas, while MCs send projections to more distributed and integrative targets such as the piriform cortex, cortical amygdala, and entorhinal cortex^1,2,57^.

### Heterogeneity in Sister Cell Responses

Although tuning profiles among sister cells were overall conserved, sister cells showed distinct differences, with temporal response properties, in particular, exhibiting marked diversity. Across approximately one-third of odour-cell pairs, response dynamics varied substantially between sister cells. This is in agreement with findings from optogenetically identified putative sister cell pairs ^62–64^, supporting the idea that temporal structure is a key axis of odour coding. Mechanistically, this diversity may arise from several factors: Firstly, differences in wiring at the level of the glomerular microcircuitry can influence how sister cells integrate excitatory and inhibitory inputs^58,59,79–83^. Secondly, lateral inhibition from granule cells (GCs) is thought to differentially modulate MCs and TCs based on their spatial positioning within the OB^84–86^. Consistent with this notion, we found that lateral dendritic geometry can in some cases predict response similarity in MCs **(Figure 5e-g)**. Finally, intrinsic cellular properties, including differences in dendritic integration and excitability, might further amplify cellular heterogeneity^87,88^.

### State- and Stimulus-dependence of Sister Cell Diversity

Our study aims to provide a foundational understanding of structure-function relationships in the OB at scale. In order to minimise the influence of top-down feedback and behavioural modulation, we therefore performed functional imaging experiments in anaesthetized mice. Differential centrifugal innervation of both glomerular and granule cell microcircuitry could allow for task-specific diversification of inhibitory inputs thereby increasing or decreasing differences between sister cells in a context-specific manner. M/TCs and in particular GCs are thought to be more active in engaged mice^89,90^ (see however ref ^91^) which could amplify the role of lateral inhibition in diversifying sister cells in a task-specific manner. Moreover, we used a panel of 47 monomolecular odours to cover chemical space. Even larger panels of odours or odours at different concentrations^92^ might further disentangle glomerular similarity. Multi-component mixtures activating the OB in a more widespread manner might further uncover differences in MC responses: Given the strong feedforward inhibition they receive, MCs may utilize their longer integration timescales to differentially integrate information across the OB and extract higher-order odour features. Dynamic stimuli^93–97^ in turn are likely to differentially engage the glomerular microcircuitry, thereby providing another substrate for diversifying sister cell activity.

### X-ray Imaging: Large-Scale Structure-Function Analysis

Here, for the first time, we combined functional imaging with synchrotron nano-holotomography to reconstruct anatomy at the mm³ scale at subcellular resolution. While this resolution was sufficient to delineate fine dendritic structure for M/TCs reliably, it did not allow identification of synapses or other connectomic features such as axons or spines. As X-ray tomography leaves the sample intact, subsequent targeted volume EM (vEM) will allow delineation of the M/TC-GC connectivity (or glomerular structure) in smaller subvolumes amenable to routine vEM approaches. Importantly, we can trace any dendrites in these subvolumes back into the larger XNH (or even larger SXRT) volumes, allowing functional annotation for the subset of imaged M/TCs. Several (9+) subvolumes could be acquired from an individual sample using recently introduced femtosecond laser milling technology^68^.

To acquire connectomic information at scale, the resolution of X-ray tomography has to be further improved. X-ray ptychography has been shown to yield resolutions at the sub 5 nm level^51^, and new radiation-hard materials have already allowed ptychographic tomography imaging on brain samples at near-synaptic resolution^9^. Notably, for the full-field XNH approach, different algorithms have recently been proposed to improve resolution through, e.g. probe retrieval^98^ or coded apertures^99^. Moreover, with the development of new dedicated nano-bioimaging beamlines, these can be optimised for <20 nm resolution with improved mechanical stability and beam focus and for higher efficiency direct X-ray detectors. Together, this promises to augment the reliability and scalability of XNH and extend the resolution to the connectomics level. Importantly, synchrotron X-ray imaging is designed to give global users access and support at no cost to the user, therefore promising a widely accessible entrypoint into connectomics.

## Conclusion

We have established a powerful correlative imaging pipeline that integrates functional information *in vivo* with structural data at subcellular resolution across mm³-scale brain volumes. This workflow’s robustness allowed us to replicate findings across three biological replicates, enabling structure-function analysis in the OB at scale. This allowed us to show that sister cells preserve glomerular identity while efficiently encoding odours in diversified response profiles.

As X-ray tomography technology continues to evolve, the framework established here opens new avenues for linking structure and function across increasingly complex neural circuits. Future developments will not only make such approaches more accessible but also enable structure-function analyses at the brain-wide scale, paving the way for mechanistic investigations into how neural circuits transform sensory input into perception and behaviour.

## Materials & Methods

### Animals

We collected samples Y391 and Y489 from animals of the following crossing: MOR174/9- eGFP^1,3^ crossed into a Tbet-cre driver line^100^ (JAX stock #024507) crossed with a GCaMP6f reporter line ^93,101^ (JAX stock #028865). Sample C432 contained an additional crossing with M72-IRES-ChR2-YFP^102^ (JAX stock #021206) **(Supp Table 2)**. All animal protocols were approved by the Animal Welfare and Ethical Review Body of the Francis Crick Institute and the United Kingdom Home Office under the Animals (Scientific Procedures) Act 1986 under licenses PA2F6DA12 and PP0015888.

### Surgeries

We anaesthetised animals with an intraperitoneal injection of sleep mix (0.05 mg/kg fentanyl, 5 mg/kg midazolam, 0.5 mg/kg medetomidine). Following shaving, we disinfected the skin over the skull with 1% chlorhexidine. After skin incision, a custom head-fix implant was fixed to the skull with medical super glue (Vetbond, 3M, Maplewood MN, USA) and dental cement (Paladur, Heraeus Kulzer GmbH, Hanau, Germany; Simplex Rapid Liquid, 437 Associated Dental Products Ltd., Swindon, UK). Next, we exposed the skull above the olfactory bulbs through a central hole in the implant and made a ∼2 mm-diameter craniotomy over the left bulb with a dental drill (Success 40, Osada, Tokyo, Japan). Throughout the procedure, we moisturised skull and brain with ACSF (10 mM HEPES, 10 mM glucose, 125 mM NaCl, 5 mM KCl, 2 mM MgSO4.7H2O, 2 mM CaCl2.2H2O, pH = 7.4). After removal of the exposed dura, we applied a drop of warm 2% low-melt agarose diluted in ACSF onto the brain, quickly followed by the placement of a 2.5 mm-diameter glass coverslip (borosilicate glass #1 thickness (150 μm)). We fixed the coverslip onto the skull with medical super glue. For samples Y391 and Y489, we injected the animals with 20 μL of dexamethasone 2-4h before surgery.

### Odour delivery

We delivered a panel of 47-50 monomolecular odourants **(Supp Table 1)** (Sigma-Aldrich, St. Louis MO, USA) through a series of custom-made 6-channel air-dilution olfactometers^93,103^ **(Supp Fig 1_1a)**. Briefly, each olfactometer consisted of a 15 ml odour bottle ((27160-U, Sigma-Aldrich, St. Louis MO, USA) containing 2ml of pure odorants, sandwiched by a pair of solenoid valves (VDW10AA, SMC). When a pair of valves opened, carrier air pushed saturated odour into the airstream, where it was diluted to 1:20, 1:9.7 and 1:4.3 for samples C432, Y391 and Y489, respectively, with a total flow rate of 1-1.5 L/min. The olfactometer was connected to a short tube positioned <1 cm away from the ipsilateral nostril to the imaged bulb. Respiration was monitored with a flow sensor (A3100, Honeywell, NC, USA) at the nostril contralateral to the imaged bulb and its signal digitised by a Power 1401 ADC board (CED, Cambridge, UK) as described before^59^. Each odour presentation was a 2 s-long square pulse quantified with a photoionisation detector (200B miniPID, Aurora Scientific) **(Supp Fig 1_1b)** and was triggered by inhalation for samples Y391 and Y489 **(Supp Fig 1_1c)** using spike2 (CED, UK). An inter-stimulus interval of >41.8 s was used to flush clean the airstream. Within each olfactometer, we randomised the order of odour presentations and repeated the entire panel two to four times per animal. We controlled the olfactometer with the custom software PulseBoy^93,97^ (https://github.com/RoboDoig/PulseBoy).

### In vivo 2-photon imaging

After surgery, we head-fixed the anaesthetised mice under the objective of a 2P microscope (Scientifica Multiphoton VivoScope). We imaged the activity of olfactory bulb neurons with 910-940 nm light from a MaiTai DeepSee laser (Spectra-Physics, Santa Clara, CA) using a 16x 0.8 NA water-immersion objective (Nikon). 512x512-pixel images were taken with a resonant scanner and a galvanometer mirror at a 30 Hz frame rate. A piezo motor (PI Instruments, UK) coupled to the objective allowed for movement along the depth axis. The FOVs for sample C432, Y391 and Y489 were 550 x 550, 330 x 730, 544 x 544 μm^2^, respectively. The total scan depths were 300-340 μm, divided into 6 planes for sample C432 and 12 planes for samples Y391 and Y489, resulting in a 5 or 2.5 Hz volume rate. We acquired samples C432 and Y391 with a constant 30 or 24 mW across depths using software SciScan (Scientifica) and sample Y489 with power increasing by depth (23-43 mW) using software ScanImage (MBF Bioscience) and gated GaAsP PMTs (Hamamatsu H11706P-40). Immediately after functional 2P imaging, the animals received an intraperitoneal injection of 50 μL of sulforhodamine 101 (0.1M, dissolved in injectable water) (Sigma Aldrich). After ∼5 min, we took an anatomical stack covering the same FOV with 1536 x 1536 pixels per image and 5 μm z-spacing with the same light and power settings. Resting GCaMP and blood vessel signals were collected in green and red, respectively **(Figure 1c)**. We used this anatomical dataset to map functionally imaged cells in XNH datasets (see *Sister cell identification and dendrite tracing*).

### Dissection and slicing

After *in vivo* 2P data collection, we sacrificed the animals, and dissected out their olfactory bulbs. Dorsal slices of the imaged bulbs were cut with a vibratome (Leica VT 1200S) into 0.6, 0.5 and 0.5 mm thicknesses for samples C432, Y391 and Y489, respectively. The 2P-imaged regions were located and included within the slice by locating the GFP-tagged glomerulus using a stereoscope (Leica MZ10 F) mounted on top of the vibratome. We used ice-cold dissection solution (4.6% sucrose, 0.563 mM CaCl2, 65.2 mM NaH2PO4.H2O, with 0.02% sodium azide, bubbled with 95% and O2/5% CO2, osmolarity 280–320 mOsm/l) to moisten and keep the tissue cold during dissection and slicing.

### Fixation and epifluorescence imaging

Next, we placed the collected slices in ice-cold fixative (2 mM CaCl2 and 0.02% sodium azide in 0.15M sodium cacodylate, osmolarity 280–320 mOsm/l, with 1% GA for sample C432 or a mixture of 1.25% GA and 2.5% PFA for samples Y391 and Y489). After 10 - 60 min of contact with the fixative, slices were briefly imaged with a widefield fluorescence microscope (Olympus Axioplan2 with LEJ “ebq 100 isolated-z” lamp). This epifluorescence dataset featured the location of the GFP-tagged glomerulus in relation to surface blood vessels and sample boundaries. It provided an estimate of the 2P-imaged region in the LXRT dataset **(Supp Fig 2_2b)**. The slices were then post-fixed overnight at 4 °C.

### Ex vivo 2-photon imaging

After overnight post-fixation, we washed the slices with 0.15 M sodium cacodylate buffer (NCB) (20-60 min each, 3 times). We then imaged the fixed tissue slices with the same 2P microscope at 800 nm excitation wavelength. In total, we acquired a volume of ∼ 0.5 x 0.9 x 0.3 mm^3^ around the GFP-tagged glomerulus with a voxel size of 0.645-1.074 μm in x, y and 5 μm in z. This *ex vivo* dataset featured blood vessels and sample surface features around the GFP-tagged glomerulus and was used to locate the GFP-tagged glomerulus in the laboratory-based micro-CT (LXRT) dataset.

### Staining and embedding

For staining, we then processed the samples with a modified heavy metal staining protocol based on^73^: 2% OsO4 in 0.15M NCB (1.5 h, 20 °C) followed by 2.5% potassium ferrocyanide in 0.15M NCB (1.5 h, 20 °C) without water wash; 1% thiocarbohydrazide (aq., 45 min, 30 °C); 2% OsO4 (aq., 3h, 20 °C); 1% uranyl acetate (aq.) warmed to 50 °C for 2 h after overnight at 4 °C; lead aspartate (aq., pH = 5.0, 2 h, 50 °C) with water washes between each step. Samples were then dehydrated with 75%, 90%, 100%, and 100% ethanol, transferred to propylene oxide and infiltrated with 25%, 50%, 75%, 100%, and 100% hard epon diluted in propylene oxide, and polymerised for 72 h at 60–70 °C.

### Laboratory-based micro-CT (LXRT)

For quality control and to allow further targeting, we imaged stained and embedded samples with a Zeiss Xradia 510 at 40 kV and 3W with LE2, LE3 or LE4 filters, achieving a 10-50% transmission. 1601 projections were taken over 180° of sample rotation with 3-20 s exposure time per frame, giving ∼ 5000 photon counts per pixel. Pixel sizes ranged from 3-6 μm. We used LXRT datasets to screen for macroscopic staining artefacts **(Supp Fig 1_8)** and to guide sample trimming **(Supp Fig 1_3)**.

### Synchrotron X-ray computed tomography with propagation-based phase contrast (SXRT)

After LXRT, samples were imaged at the TOMCAT beamline at the Swiss Light Source (Viligen, Switzerland) **(Supp Fig 1_2)** to verify staining integrity at the subcellular level and to allow precise targeting for XNH. A multilayer monochromatic X-ray beam was filtered with 0.1 mm-thick aluminium and 0.01 mm-thick iron to 21 keV before reaching the sample, placed 50 mm before the detector. The detector unit consisted of a scintillation screen (LuAG:Ce 20 μm) used to convert transmitted and scattered X-rays to visible light and an objective lens (20x Olympus UPlan Apo) coupled with a CMOS camera (pco edge 5.5) used to focus and digitise the visible light photons. The sample was rotated by 180 degrees via 2601 steps, with a projection image taken at each step with 300 ms exposure time. Each scan took ∼13 min and gave an 832 x 832 x 702 μm^3^ volume with an isotropic voxel size of 325 nm. Each sample was tiled with ∼20 scans with 0.25 mm and 0.03 mm overlap in x and y, respectively, resulting in a total acquisition time of 3-4 h per sample.

We reconstructed each scan individually with the ‘GridRec’ pipeline^104^. For phase retrieval we set the refractive index parameters to δ = 2e-6 and β = 6e-7, and applied an unsharp mask filter with a stabiliser of 0.6 and a Gaussian kernel filter of 1.0. We then stitched all reconstructed scans into a continuous volume with a 3D non-rigid local pairwise stitching algorithm NRStitcher^105^.

Sample C432 was also imaged at the I13-2 beamline at Diamond Light Source (Didcot, UK) as described before^6^. The dataset was of similar quality and equivalent in its use to the one collected at TOMCAT.

### Sample trimming

To meet the sample size requirement of XNH while preserving the entire 2P-imaged regions we then trimmed the samples. To do this, we warped the 2P FOV into the SXRT dataset, which was then warped into the LXRT dataset^106^. We planned trim lines on 3D renders of the LXRT dataset in Amira (Thermo Fischer Scientific). Samples C432 and Y391 were trimmed into oblique prisms with an ultramicrotome (Leica UC7) loaded with glass and ‘trim 90’ diamond (Diatome) knives in two to three iterations. An LXRT dataset was acquired, and trim lines were redesigned after each iteration. Sample Y489 was trimmed into a ∼1 mm diameter cylindrical pillar with fs-laser milling^68^ **(Supp Fig 1_3)** at Carl Zeiss Microscopy GmbH in Oberkochen, Germany. The pillar geometry prevented streak artefacts that arise from sharp edges^107^. To mill and monitor the sample we used a Zeiss Crossbeam 350 FIB-SEM microscope. Firstly, we ablated unnecessary regions of the sample (laser time ∼5 min). Secondly, we polished the sample surface using a concentric ring pattern (laser time ∼2 min). The femtosecond laser (TruMicro 2000, TRUMPF) was tuned to 515 nm with a pulse duration of <350 fs and operated at ∼2.5 W under ∼50 mbar nitrogen pressure. SEM imaging was done at 5 kV, 200 pA with a SESI detector.

### X-ray Nano-Holotomography

After trimming, we imaged the samples at the ID16A beamline of the European Synchrotron Radiation Facility (Grenoble, France) **(Supp Fig 1_4, Supp Table 3)**. The ID16A beamline endstation is located 185 m away from the undulator source. Unlike SXRT methods, which use parallel X-ray beams, XNH uses a focused beam to achieve geometric magnification. Fixed-curvature, multilayer-coated Kirkpatrick–Baez allow to produce a nanofocus with a size of ∼30 nm at 17 keV and ∼15 nm at 33.6 keV ^52^. We placed the sample on a high-precision rotation stage^108^. The detector was composed of a 23 μm-thick Europium-doped Gadolinium Gallium Garnet (GGG:Eu) scintillator, coupled with a charged-coupled device (4096 x 4096 pixels FReLon for sample C432 or 6000 x 6000 pixels Ximea for samples Y391 and Y489, both used with on-chip binning to 2000 x 2000 pixels)^109^ placed 1.2 m downstream of the sample. Both the X-ray optics and the sample stage were located inside a vacuum chamber, kept at ∼10^-8^ mbar, while the detector unit was placed outside the chamber at ambient pressure.

We imaged the sample at four propagation distances by changing the sample position between the focal spot and the detector. At each location, the sample rotated through 180 degrees either step-by-step or continuously, during which a fixed number of projections were taken. Each projection image was a hologram, which recorded the self-interference patterns of X-rays after they passed through the sample and were allowed to propagate **(Supp Fig 1_4a)**. We covered the entire sample volumes with up to 36 tiles **(Supp Fig 1_4b)**. Note that the sample was acquired at each location (i.e. tile) with all distances before moving to the next location.

Samples C432 and Y391 were acquired in ‘step-by-step’ mode. This allowed random (<25 px) lateral displacements controlled by high-precision piezoelectric actuators^53^ to be added at each rotation angle, which effectively prevented ring artefacts in the final reconstructed volume. Sample Y489 was acquired with a faster ‘continuous’ mode with a novel dose distribution method that allowed subsequent data denoising **(Supp Fig 1_4c)**. Specifically, a tile was first acquired at the planned location and then again 5 μm displaced (in x, y and z). The exposure time was halved per acquisition, so the combined exposure time (i.e. X-ray dosage) per projection was kept constant. The two acquisitions were then computationally combined (see below) to give one volume per tile location. Samples C432 and Y391 were acquired at cryogenic conditions and sample Y489 at room temperature. For cryogenic imaging, the vacuum chamber was kept at 120 K with liquid nitrogen and the sample was transferred into and out of the chamber with a Leica cryo-shuttle. The imaging parameters for all samples are listed in **Supp Table 3**. The total imaged volume and acquisition time are 0.25 mm^3^ in 91 h, 0.37 mm^3^ in 80 h and 0.40 mm^3^ in 66 h for samples C432, Y391 and Y489, respectively.

For data reconstruction, each projection image was pre-processed to correct for optics and detector-specific distortions and noises and then normalized with the empty beam projection. For each rotation angle, the four projections from the four distances were aligned by matching sample features. Phase maps were then calculated from the aligned projections using an iterative algorithm^7,110–112^. The refractive index decrement δ and the absorption index β were set to that of osmium (δ/β = 9 for X-ray energy of 17 keV, δ/β = 27 for 33 keV) and the initial amplitude and phase terms were approximated by a method adapted for multiple propagation distances^111–113^. After this initial parameter estimation, the phase term was updated while the amplitude term was held constant at each iterative step. After the phase term converged (usually < 10 iterations), phase maps were calculated for each aligned projection independently. Lastly, phase maps from all rotation angles were combined into a 3D volume using filtered back-projection^114^. In each reconstructed tile, the 200 µm diameter x 200 μm height cylindrical core captured by all distances had the highest resolution (100 nm isotropic voxels) with an edge of progressively lower resolution that extended to the full 321.6 µm diameter, 321.6 μm height cylindrical field of view.

Scans acquired in continuous mode were denoised after reconstruction using the denoising pipeline we developed recently^71^ **(Supp Fig 1_4d)**. In continuous mode, each sample location was acquired twice with a slight positional shift (see above). The two image volumes were first reconstructed independently and then aligned with the pi2 library^105^. The aligned volume pair contained the same tissue features but uncorrelated artefacts, which can be removed using the Noise2Noise algorithm^70^. For this, pairs of 3D patches were extracted from pairs of aligned volumes and used to train a 3D U-Net^115^. The network had 4 downsampling blocks in its encoder part and 56 filters for the first convolution. The patch size was set to 96x96x96 and the batch size to 32. The data augmentation process was composed of a random rotation and a random horizontal flip. The learning rate used was 5x10^−4^. Training was conducted using the Adam optimizer and the Mean Squared Error loss function. Details on the inference process are available in ref ^71^. To speed up the process, all tiles from a sample were split into three groups based on the features they contained. The features assessed were both biological e.g. glomeruli, external plexiform layer, granule cell layer, and non-biological e.g. resin or empty space. For each group, only one tile (the tile with the best alignment quality) was used to train a neural network from scratch. Once trained, each model was fine-tuned on the remaining tiles from the same group. The resulting image quality was comparable to that with a full training approach. For samples with tens of tiles like ours, this transfer learning approach saved hundreds of hours of processing time, as fine-tuning took only around one hour while full training took dozens of hours. After all processing, all tiles were stitched into a continuous volume, via an implementation of NRstitcher ^105^ on webKnossos^116^.

### Glomerulus and nuclei segmentation

To segment nuclei **(Supp Fig 1_6c)**, we developed an image classifier consisting of a trained U-net followed by a heuristic filtering of false positives based on the expected compactness and volume of cell nuclei^72^. In brief, multiple (50 µm)^3^ subvolumes sampling the different histological layers contained in the XNH datasets (glomerular, external plexiform, mitral and granule cell layers) were defined as bounding boxes in webKnossos^116^. Expert annotators (Mindy Support) then accessed these subvolumes, only displaying data downscaled two-fold to (200 nm)^3^ voxels. Each annotation task involved locating all nuclei in a subvolume with single-node skeletons, followed by a manual volume segmentation of all nuclei giving each nucleus a different segment identifier. The trained classifier generated segmented volumes on each dataset. We then generated meshes on those segmented volumes^117,118^, restricting the simplification error to 16000 nm. For each segmented object, we computed its volume (number of voxels), centroid (Euclidean distance transform of all voxels), surface (sum of the areas of all faces of its mesh), and sphericity (ratio of the surface-to-volume of an object to the one of a sphere of the same volume^119^). For each run of the image classifier, a summary table was generated listing these four variables for all segmented objects. Cutoffs were identified for sphericity and volume that would retain the vast majority of nuclei while discarding non-nuclei- segmented features (not round, too small). Together, the classifier followed by the heuristic filter delivered a robust segmentation of nuclei (precision and recall > 0.9)^72^.

We segmented all glomeruli present in each dataset **(Supp Fig 1_6d)**through a consensus manual segmentation approach. A complete XNH dataset was submitted to three manual expert annotators (Mindy Support) using the webKnossos task system. Each annotator would explore the dataset downscaled 32-fold (3.2 µm voxels), only seeing a single orthogonal projection (xy, xz or yz). This provided both three-fold redundancy as well as a segmentation accuracy as close as possible to isotropic. The manual segmentation task had two steps: first, identifying all glomeruli in the volume using single-node skeletons, followed by volume segmentation of each glomerulus in the list using different segment identifiers. A final proofreading ensured all tasks had identified the vast majority of glomeruli. To find the consensus segmentation, first, we found the correspondences between segments across tracers. Then, we applied a voxelwise majority voting algorithm to calculate the shape of the segmentation object, keeping voxels with ≥2 votes and removing voxels with only 1 vote. Glomerular segments were ensured therefore to be non-overlapping. This segmentation returned all glomeruli independently segmented in 3D, with a different identifier. By matching these identifiers with a warped contour of the regions of interest delineating individual glomeruli in 2P data, we filtered the subset of glomeruli that were analysed in vivo **(Supp Fig 1_6e)**.

### 2P data processing

Each i*n vivo* 2P time series was aligned with suite2p^120^ (https://github.com/MouseLand/suite2p). For suboptimally aligned planes, we applied rigid transformations extracted from well-aligned planes using a custom script. After registration, cellular and glomeruli ROIs were automatically detected and drawn in suite2p, respectively, and their odour responses were extracted **(Supp Fig 1_9)**.

### Sister cell identification and dendrite tracing

We first mapped 2P ROIs in the anatomical dataset (see *In vivo 2-photon imaging*), which was warped to the XNH datasets using BigWarp^106^ with manually seeded landmarks present in both datasets (e.g. blood vessel branching points, **Figure 1c**). We then built a warp graph, where nodes are individual datasets, and edges the warp relationships between a given dataset pair (paired landmarks and warp function). This graph allows combining warps across linked datasets, such as moving from anatomical 2-photon to XNH, as described previously^6,73^. We then warped the ROI seeds to the XNH dataset using the WarpAnnotations toolbox (https://github.com/FrancisCrickInstitute/warpAnnotations). Most ROI seeds were warped into unique cells in the XNH datasets. Finally, we inspected each seed manually to be assigned to the correct XNH soma using surrounding somatic and dendritic patterns.

To identify sister cells, we traced the apical dendrite of each functionally imaged ROIs in webKnossos^116^ **(Supp Fig 1_5, Supp Fig 1_6f, Supp Video 1,2)**. Each apical dendrite tracing consisted of nodes seeded within the dendrite as it was followed from the soma to its parent glomerulus and edges that connected the nodes. The vast majority of cells contained only one apical dendrite, the one cell with two apparent apical dendrites was discarded from further analyses. Sister cells were defined as cells that project their apical dendrites to the same parent glomerulus. All ROIs that were ambiguously mapped or traced were discarded from further analyses **(Supp Table 4)**. Note that some parent glomeruli were not functionally imaged (i.e. outside of 2P FOV) and non functionally imaged sister cells of a glomerulus were not included in this study.

Similar to apical dendrite tracing, lateral dendrites which consisted of branchings that extend across the external plexiform layer that originated from the soma or the apical dendrite were exhaustively traced in webKnossos for cells from three glomeruli **(Supp Fig 5_4, 5_5, 5_6)**.

### Circuit delineation and rendering

We traced glomerular columns imaged *in vivo* based on XNH data tracings **(Supp Fig 1_6b)**. Apical dendrites were associated with a glomerulus if the node closest to the glomerular layer was on a voxel belonging to that glomerulus. Similarly, apical dendrites were associated with the automatically segmented nucleus based on the coordinates of the closest node of the dendrite to the mitral cell layer. This allowed grouping objects as cells (nucleus + apical dendrite) and ultimately delineated the excitatory backbone of the glomerular columns imaged *in vivo* (glomeruli + sister cells shown in the same colour). Volume renders of datasets and segmented features were generated using SideFx Houdini 19.5.303 and Maxon Redshift 3.5.8 **(Supp Videos 3,4)**. Datasets were exported from webKnossos as image stacks using its Python API. Segmented nuclei were converted into a polygonal mesh using the tools described in ref ^121^.

### Odours in chemical space

Chemical properties of 5103 chemicals (both odorants and odourless, from ref ^122^) were collected from the Modred descriptor database^123^, using the RDKit toolkit (https://github.com/rdkit/rdkit). For each chemical pair, we calculated the Canberra distance of all descriptors. For visual display, we created a two-dimensional embedding of the chemical space using multidimensional scaling of the pairwise distances. The embedding of the stimulus set used for sample Y489 is shown in **Supp Fig 1_7**.

### Sister cell analyses

#### Odour responses

The odour response of each cell-odour pair was defined as the z-score of F, averaged across all repeats (n = 4 for samples Y391, Y489; n = 2 for sample C432). Analogously to previous work^124–127^, we defined the average dendritic signal of all Tbet+ cells in the glomerulus as the “glomerular odour response”. The comparison of parent glomerular signals to sister cell signals is shown in **Supp Fig 3_2** and confirms that the glomerular signal is indeed close to the average signal of M/TCs (provided a sufficient number of sister M/TCs is imaged simultaneously).

For most analyses, a strict criterion was used to subset cells with evident odour responses to minimize the effect of any imaging noise in the conclusions of this study. A cell was considered “strictly responsive” if its odour response (i.e. repeat-averaged z-score of F) was > 3 or < -3 for at least 1 s consecutively, for at least 1 odour within 22 s after odour onset.

#### Response integrals and tuning correlations

The response integral of each cell-odour pair was defined as the integral of odour response over the odour presentation period (t = 0-2 s) apart from the following exception. With the experimental volume rate, the standard deviation of baseline fluorescence was estimated using ∼ 8 data points which could potentially introduce spurious fluctuations in response amplitude. For a more accurate estimate of trial-to-trial variability and an assessment of how sister correlation compares to it, the boxplots in **Figure 2c1** were made using z-score derived with global baseline standard deviation (i.e. estimate from the entire baseline trace per cell), and uses t = 0-7 s response integral. Results with usual z-score over a range of integral window sizes are available in **Supp Fig 2_1 d**. No matter the metric, the conclusion that sister cell tuning correlation is similar to self correlation across repeats always holds.

The odour tuning of a cell was defined as the vector of its response integrals for all odours. Correspondingly, we defined odour tuning correlation between two cells as the Pearson correlation value of their odour-tuning vectors. Glomerulus-cell tuning correlation was similarity defined between a glomerulus and a cell.

To assess how reliably a cell responded to odours across repeats, we used a split-repeat correlation, defined as the Pearson correlation value of a cell’s tuning calculated with responses from repeats 1, 3 versus the tuning calculated with repeats 2, 4. A cell that responded faithfully would have a high split-repeat correlation and a cell that responded to some but not all repeats a low value. We calculated split-repeat correlations for each individual cell that acted as a reference value to judge sister cell similarity.

Tuning correlation boxplots were drawn so that the box covered the first and third quartile of the data, the midline represented the median, and the whiskers extended to the most extreme, non outlier data points. Outliers were defined as data points that were either smaller than the first quartile - 1.5 times the interquartile range or larger than the third quartile + 1.5 times the interquartile range.

#### Strong odour overlap and weak odour correlation

In **Figure 3** we introduce the “strong-odour overlap” as a metric to assess the identity of the strongest activating odours across sister cells. For each cell, the top n (e.g. 5) strongest activating odours were identified by ranking the response integrals across all odours. We defined “strong-odour overlap” as the fraction of shared strong odours per cell pair. A fraction of 1 implied that the top n strongest activating odours were completely shared between a pair of cells. A fraction of 0 implied that the cell pair responded to completely different odours most strongly. In the absence of any structure or similarity between sister cells (null hypothesis), the fraction of overlap increases with increasing n. E.g. When n equals the total number of odours in the stimulus panel, the overlap fraction will always be 1. Therefore, sister and non-sister strong-odour overlap was plotted against chance level, which was simulated by randomly sampling two independent sets of n odours 1000 times. Additionally, strong-odour overlap was also calculated for each individual cell using split-repeat odour responses (i.e. averaging repeat 1, 3 versus 2, 4), which like split-repeat tuning correlation, reflected odour response reliability of cells.

To assess the similarity of sister cell responses to weak odours, glomerulus-cell tuning correlation was calculated with n strongest odours removed. Glomerulus tuning acted as an independent reference that sister cell tunings can be compared to. The cell-cell correlation equivalent is shown in **Supp Fig 3_1**.

#### Glomerular and cellular signal comparison

The glomerular signals measured in this study reflect the average dendritic signals from all Tbet+ sister cells per glomerulus. Glomeruli signals were extracted from manually drawn oval ROIs **(Supp Fig 1_9)**. ROIs were consolidated with glomeruli segmentation in the XNH dataset to avoid splits or mergers^6^. Signals from glomeruli sampled in more than one plane were averaged together. To understand the relationship between glomerular and cellular signals **(Supp Fig 3_2)**, the tuning correlation was calculated between the glomerular signal and the mean odour responses of sister cells using increasing numbers of sisters. To avoid the potential confounder of some sister cells being more or less similar to the parent glomerulus, for n sister cells, we randomly sampled n sister cells 50 times, calculated the mean odour responses of each sample, and correlated this to the parent glomerulus response vector. In the figure, we plotted the mean and the 95% confidence interval of these correlation values.

#### Response shape and temporal similarity index

In **Figure 4** and in the associated supplementary figure **Supp Fig 4_1** we introduce the “response shape” and the “trace-similarity index” to compare the temporal structure of sister cell odour responses. Response shape was defined as the z-score of F normalised by the standard deviation of the trace itself and highlighted differences in sister cell responses that would otherwise be hard to spot amid varying response amplitudes **(Supp Fig 4_1)**.

The temporal-similarity index is a metric that measures the average similarity of response shapes across sister cells per glomerulus-odour pair. Firstly, odours that reliably elicited responses in at least one sister cell were identified for each glomerulus (using the aforementioned strictly responding criterion). A response was considered reliable for a cell-odour combination if the average z-score correlation across all repeat pairs was >0.8. For these odours, we calculated the Pearson correlation of odour responses (i.e. repeat-averaged z-score of F) for every sister cell pair and defined its mean as the temporal-similarity index per glomerulus-odour pair. An arbitrary threshold of < 0.7 was considered “diverse” and ≥ 0.9 “stereotypical”

#### Glomerulus, cell and odour decoding

To assess the level of response dissimilarity among sister cells, a support vector machine (SVM) with a linear kernel was used to predict individual cell identity. The classifier was trained on cellular response integrals calculated from a random sample of three repeats per cell-odour pair with cell identity as the training label. The classifier was tested on the remaining repeat. If sister cells were very similar, the classifier would be expected to confuse the identity of a cell with its sisters. This can be quantified as ‘glomerulus prediction accuracy’, where any prediction within the parent glomerulus of a cell was considered a hit and prediction to non-parent glomeruli a miss.

To assess the function of sister cells in odour decoding, a different linear SVM was trained to predict odour identity using cellular response integrals, as a function of increasing number of sister cells **(Supp Fig 4_2b)**. To remove the confounder of increasing olfactory receptor coverage, the first data point included exactly one sister cell from all glomeruli. As sister cell n increases, up to n sister cells per glomerulus were included in the classifier. For glomeruli with < n total sister cells, all of its sisters were included. As a reference, the effect of increasing olfactory receptor coverage was demonstrated by plotting odour prediction accuracy against increasing number of glomeruli **(Supp Fig 4_2c)**, using a similar linear SVM approach.

#### TC vs MC and the effect of denoising

Projection neurons were categorized into TCs and MCs by soma depths. Further division of TCs into external or middle TCs based on lateral dendrite number did not show significant differences and hence they were grouped together. Sample Y489 contained both TCs and MCs while sample Y391 contained predominantly TCs and C432 MCs.

Imaging deeper typically results in reduced signal-to-noise. Thus, to ensure a fair comparison of TC and MC activity, we assessed the influence of noise on activity metrics. Odour responses were denoised with non-negative matrix factorization (NMF, the entire z-score trace was raised above 0, fitted with NMF, then brought down to baseline) with varying degrees of smoothness **(Supp Fig 5_1b)**. NMF was chosen over low-pass filtering and PCA because of their tendency to give false activity bumps at either end of the response traces. To assess the robustness of response integral with respect to noise, glomerulus-cell tuning correlation was calculated using response integrals from varying levels of denoising **(Supp Fig 5_1a)**. TC-MC tuning correlation was calculated using response integrals of TC-MC pairs **(Supp Fig 5_1d,f)**.

#### Lateral adjacency score and distances

Lateral dendrites of M/TCs sample distinct parts of the EPL where they can contact GCs. We reasoned that for two M/TCs close proximity of their lateral dendrites increases the likelihood that they contact the same GC. We thus devised the lateral adjacency score **(Figure 5e** and **Supp Fig 5_2)** as a metric to quantify this proximity (and thus the likelihood of engaging each other through the GC network). Firstly, the XNH dataset was divided into 50x50x50 μm gridlets. Then for each cell, the gridlets that its lateral dendrite occupied were listed and for a cell pair, the gridlets that contain lateral dendrites from both cells were listed. Lateral adjacency score was defined as the number of co-occupying gridlets divided by the total number of gridlets occupied by the cell pair. 50 µm was chosen reflecting an approximate width of a typical GC dendritic tree^86^.

The centroids of glomerulus contour and the location of soma nodes in XNH were used to calculate Euclidean distances between glomerulus pairs, cell pairs and glomerulus-cell pairs.

#### Sister cell models

To assess what could be the advantage of the observed similar but non-identical sister cell responses, we investigated the performance of three models of varying levels of sister cell response similarity, from the angles of odour encoding capacity and odour generalization ability.

Sister cell responses were modelled as nonlinear transformations of cell inputs, which in turn, were modelled as the parent glomerulus response plus random Gaussian noise. Glomerular odour responses were values drawn from a Gaussian distribution. Cell input matrix *Y* and cell output matrix *X* were such that their singular value decompositions follow *X = URV^T^*, *Y = USV^T^* and the relationship of their singular value matrices follows , where β controls the degree of sister cell similarity. This relationship derives from the minimization of a loss function that balances cell input-output similarity and the volume of odour representations in cell space (see **Appendix**). β = 30,000, 30, 0.003 were used for model ‘relay’, model ‘balanced diversity’ and model ‘network’, respectively, with decreasing sister cell similarity. In total, 802 cells from 20 glomeruli and 3209 odours from 10 categories were simulated. Odour categories represent perceptually similar odour classes, such as the smell of different citrus fruits or different feline species. Odours belonging to the same category were generated with the same response vector plus random values drawn from a Gaussian distribution.

Odour generalization ability of each model was quantified by a metric that measures odour category separation in cell representation space. Cell representation space was defined by a subset of randomly sampled cells. Odour category separation was visualized by PCA projection of odour response vectors onto the top two principal components of this cell representation space, and was quantified by the average silhouette score of this projection **(Supp Fig 5_7d,f)**. High inter-cluster distance (i.e. cluster separation) and low intra-cluster distance (i.e. cluster tightness) results in a high metric value.

Odour encoding capacity of each model was quantified by the odour decoding accuracy of a linear SVM which measures how linearly separable odours are in cell representation space i.e. odour discrimination ability. Odours that overlap in their representation are harder to linearly separate than odours that do not overlap. Such ‘noise clouds’ were visualized by projecting odour repeats (n=500) of three odours to a 2D plane, defined by the mean response vector of the three odours **(Supp Fig 5_7e,g)**.

## Data availability

Source data of the plots presented in the main figures are provided as a Source Data file. The datasets and major annotations reported in this study are accessible through the associated code repository (see Code Availability).

## Code availability

Analysis code and skeletal annotations are available from https://github.com/yz22015/sister_2p_xnh and curated calcium traces are available from https://zenodo.org/records/15150149.

X- ray tomography reconstruction and denoising code is available from https://gitlab.esrf.fr/tomotools/nabu and https://github.com/xni-esrf/SSD_3D, respectively.

For the three correlative experiments (C432, Y391 and Y489), all anatomical 2-photon, synchrotron X-ray phase contrast, synchrotron X-ray holotomography datasets and lab-source CT datasets are available from https://github.com/FrancisCrickInstitute/warpAnnotations/tree/main/warping/data.

The warping landmarks and functions that allow annotations to be warped across datasets are accessible on the release v1.1.0 of warpAnnotations: https://github.com/FrancisCrickInstitute/warpAnnotations

## Author contributions

YZ, CB, TA, AP, ATS conceived the project and designed experiments. YZ, CB, TA, AB, CW, PC, AP performed experiments and acquired data, YZ, CB, TA, AL, AB, JL, AN, NR, PC, AP processed and analysed data, YZ, CB, AL, AB, MB, AN, NR, AP contributed tools, YZ, ST, MK performed simulations, YZ, CB prepared figures, YZ, CB, ATS prepared the initial manuscript draft, YZ, ATS wrote the manuscript with input from all authors. CB, AB, NR, PC, AP, ATS supervised the project and beamtime experiments.

## Acknowledgements

The authors are grateful to the biological research, scientific computing, and electron microscopy science technology platforms of the Francis Crick Institute. This work was primarily supported by a Physics of Life grant (EP/W024292/1) to A.T.S. and A.P. funded by EPSRC and Wellcome. This research was funded in part by the Wellcome Trust (FC001153, 110174/Z/15/Z to A.T.S.). For the purpose of Open Access, the author has applied a CC BY public copyright licence to any Author Accepted Manuscript version arising from this submission. We acknowledge beamtimes ls2918, ls3025, ls3186, ls3231 at the European Synchrotron Radiation Facility, Grenoble, France (ESRF) and beamtimes e18026, e19556, e20104 at the Swiss Light Source, Paul Scherrer Institut, Villigen, Switzerland (PSI), as well as beamtime MT20274 at the Diamond light source, Didcot, UK. This work was supported by the Francis Crick Institute, which receives its core funding from Cancer Research UK (FC001153 to A.T.S.), the UK Medical Research Council (FC001153 to A.T.S.), and the Wellcome Trust (FC001153 to A.T.S.). A.P. acknowledges funding from the European Research Council under the European Union’s Horizon 2020 Research and Innovation Programme (852455). We thank Malte Storm (Diamond Lightsource / Helmholtz-Zentrum), Liz Duke (Diamond Lightsource / EMBL), Marie-Christine Zdora (Diamond Lightsource / PSI / Monash University) for fruitful discussions and help in particular with initial beamtime at the Diamond Lightsource, and all members of the Schaefer and Pacureanu labs, of ID16A and Adrian Wanner, Ana Diaz, Manuel Guizar-Sicairos, Tomas Aidukas, Nick Phillips, Chris Jacobsen, Aaron Kuan, and Wei-Chung Allen Lee for insightful discussions around X-ray connectomics, and Jeroen Claus, Ethan MacKenzie and the team at phospho bioimaging for 3D rendering support. Moreover, we thank Alexander Fleischmann, Florin Albeanu, Dima Rinberg, Kevin Franks, and Aaron Kuan for detailed comments on earlier versions of this manuscript.

