## Supplementary Information for "Structure-Function Mapping of Olfactory Bulb Circuits with Synchrotron X-ray Nanotomography"

This document includes the following:

- Supplementary video links
- Supplementary figures
- Supplementary tables
- Appendix texts

### Table of content:

Supp Video 1: Real time screen recording during the tracing of an example MC. ................. 3
Supp Video 2: Real time screen recording during the tracing of an example TC. .................. 3
Supp Video 3: Fly-through of XNH dataset and rendering of glomeruli and nuclei segmentation, apical dendrite tracing and sister cell odour responses of sample Y489. . 4

### Supplementary videos:

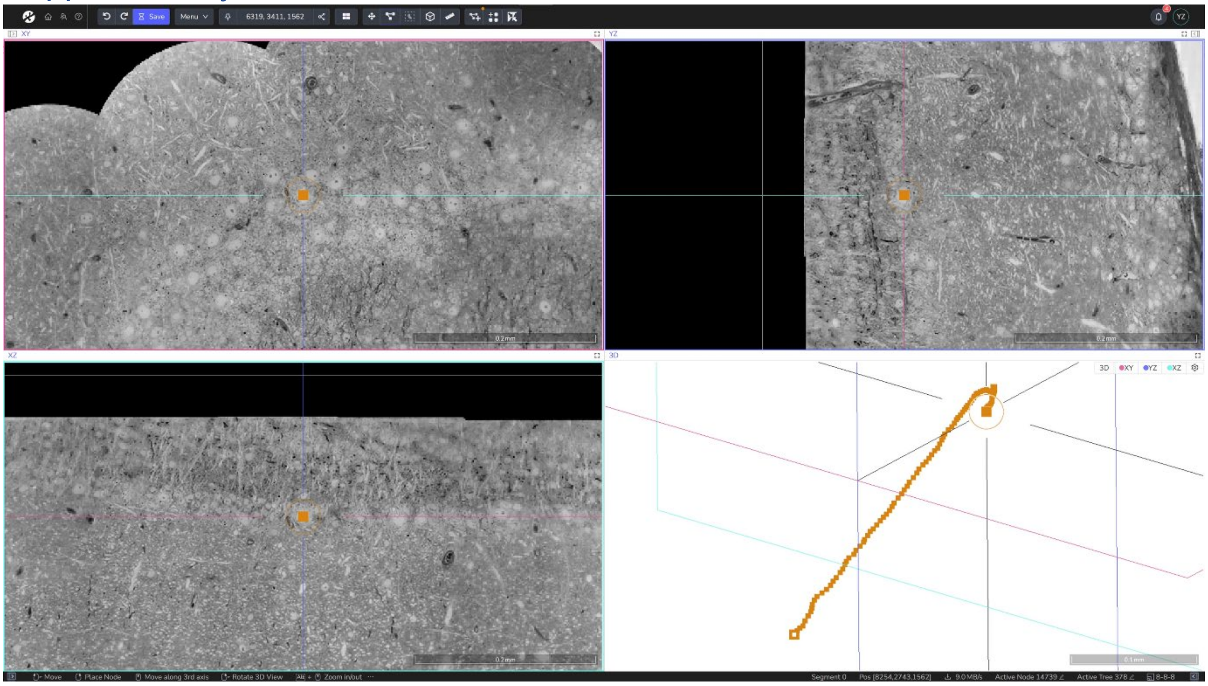

**Supp Video 1: Real time screen recording during the tracing of an example MC.**
https://youtu.be/y2n5O67MV74?si=Ct9e7LhVweo5OMdp

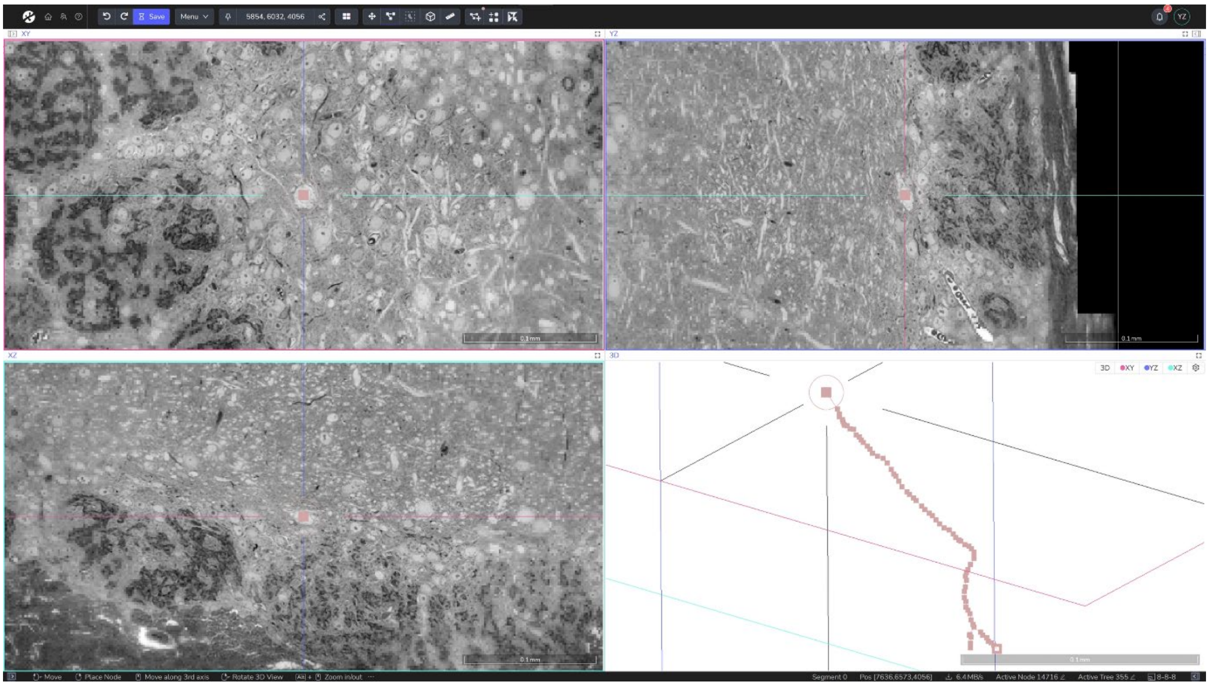

**Supp Video 2: Real time screen recording during the tracing of an example TC.**
https://youtu.be/VjkWIW1xTOo?si=7JZde2-hymDHJUZZ

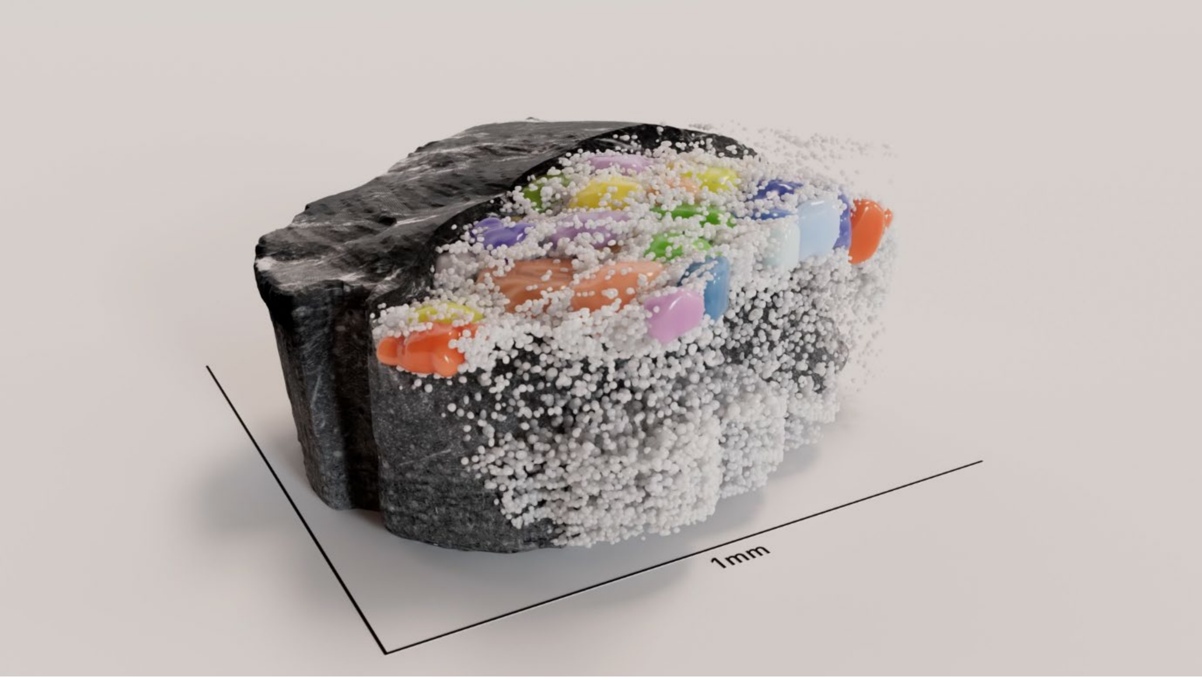

**Supp Video 3: Fly-through of XNH dataset and rendering of glomeruli and nuclei segmentation, apical dendrite tracing and sister cell odour responses of sample Y489.**
https://youtu.be/godL2-4rQKU?si=XSsPX9n_uw9cOTv6

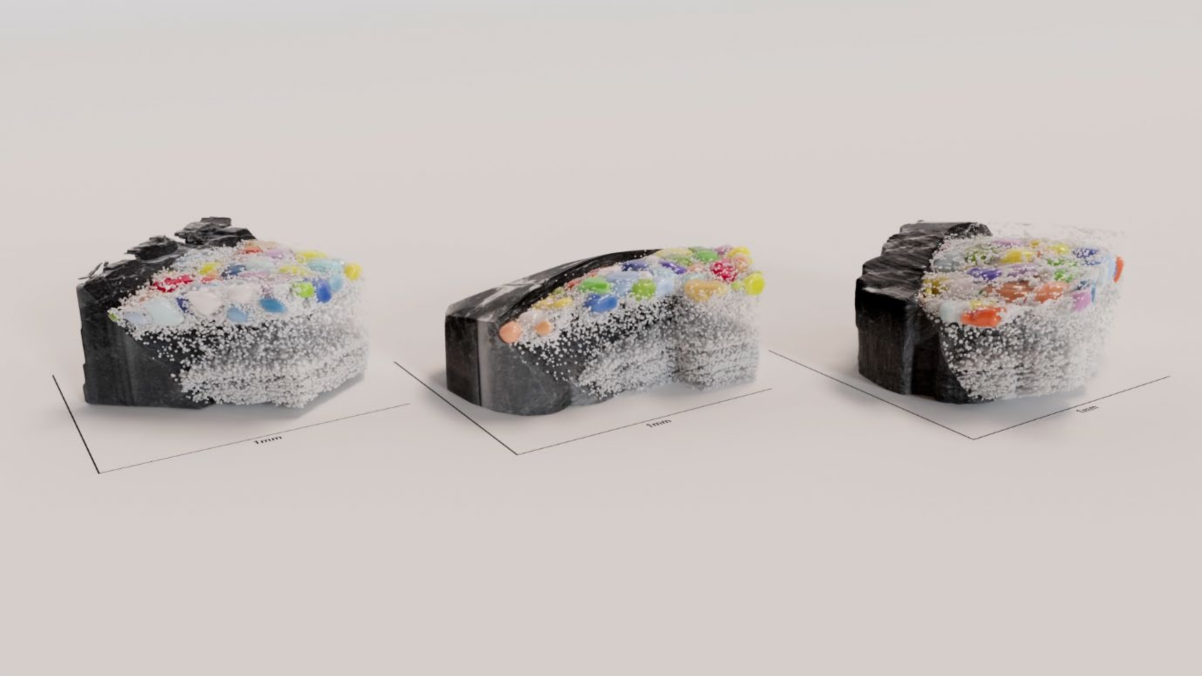

**Supp Video 4: Rendering of XNH dataset, glomeruli and nuclei segmentation, apical dendrite tracing and sister cell odour responses of samples C432, Y391 and Y489.**
https://youtu.be/ue9DysFshfQ?si=eajbgEtC2ziK5t_Y

### Supplementary figures:

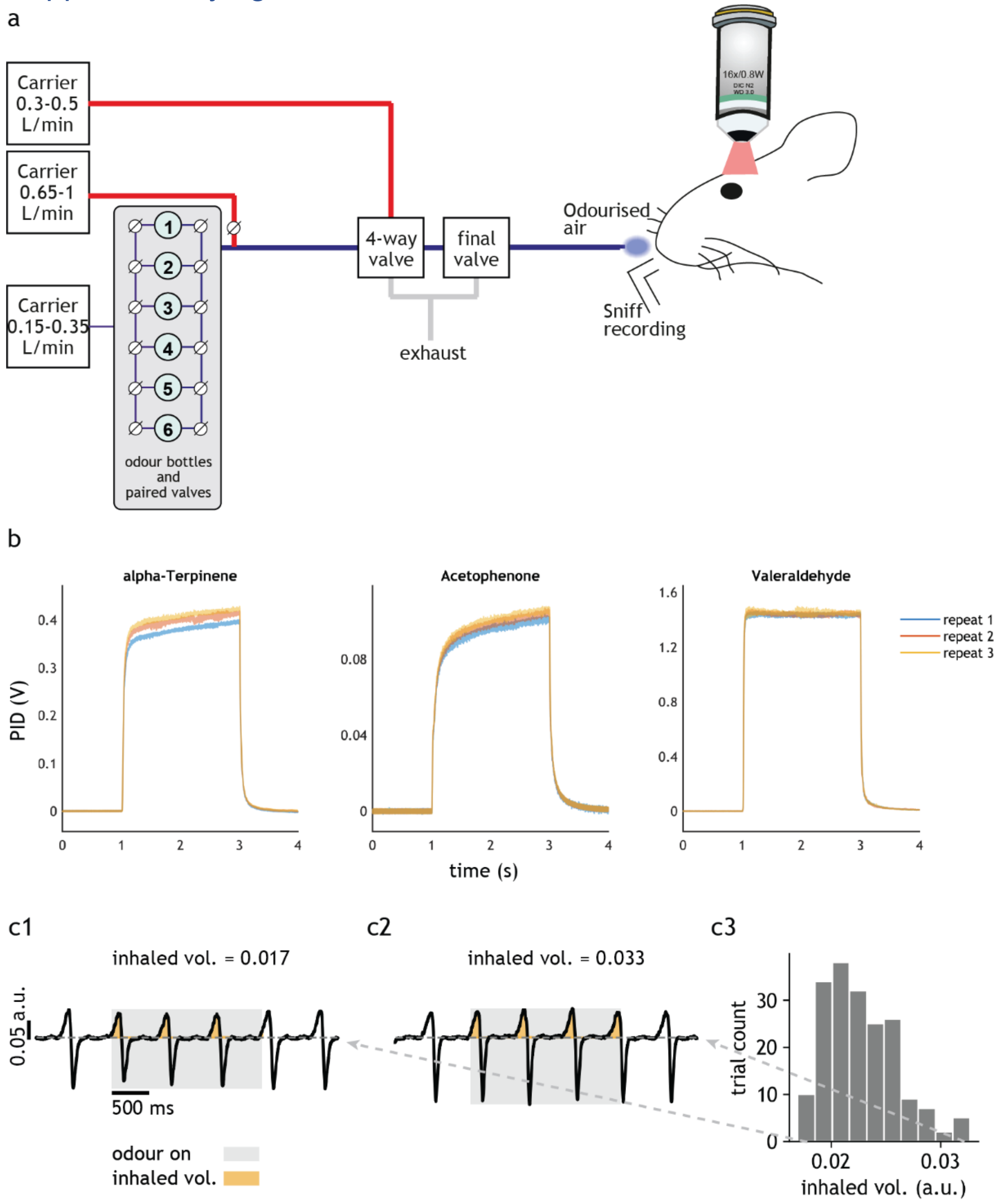

**Supp Fig 1_1: Odour delivery and stimulus shape.**
**(a)** Odour delivery device used in *in vivo* 2P imaging of mouse OB. In brief, pure odours diluted with air and controlled by solenoid valves were delivered to the ipsilateral nostril of the recorded bulb while respiration was recorded with a flow sensor at the contralateral nostril. **(b)** Shapes of 2 s-long odour pulses delivered by the odour delivery device in **(a)** of three example odours recorded with photoionisation detector (PID) on the day of *in vivo* 2P imaging (of sample Y489). Each odour was recorded three times. **(c)** Respiration traces. Odour delivery (grey shaded area) was triggered by inhalation. **(c3)** Histogram of inhalation volume of all trials from sample Y489.

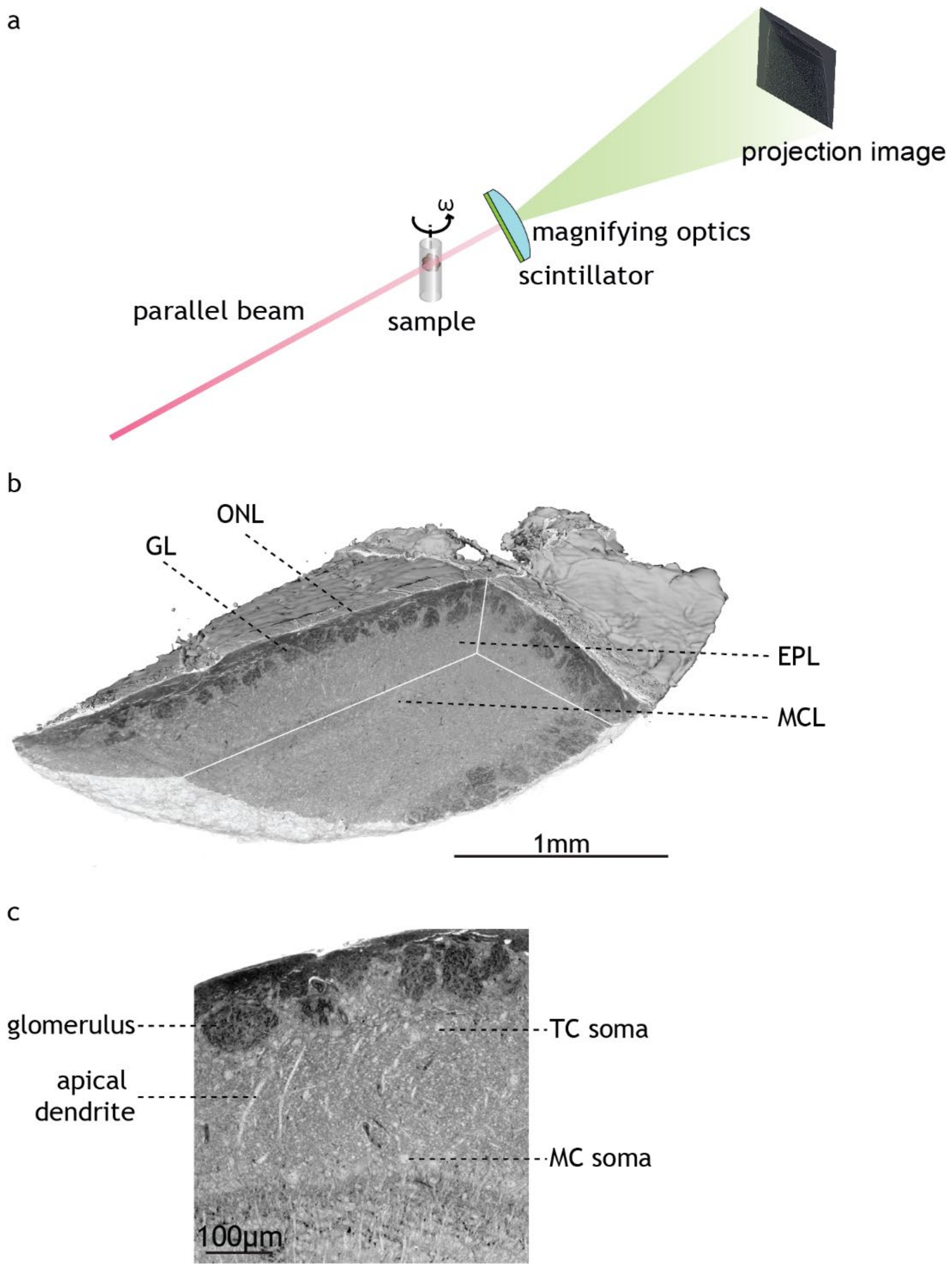

**Supp Fig 1_2: SXRT acquisition and dataset.**
**(a)** Imaging principle of SXRT. In brief, a parallel beam of polychromatic X-rays filtered to the desired energy created images of the sample while it rotated. X-ray signal was converted to and detected as visible light photons through a scintillator. **(b)** Volume render of an example SXRT dataset (sample Y489) with a cutout showcasing OB tissue layers. **(c)** Example SXRT image crop with identifiable features relevant to this study indicated. ONL, olfactory nerve layer; GL, glomerular layer; EPL, external plexiform layer; MCL, mitral cell layer; TC, tufted cell; MC, mitral cell.

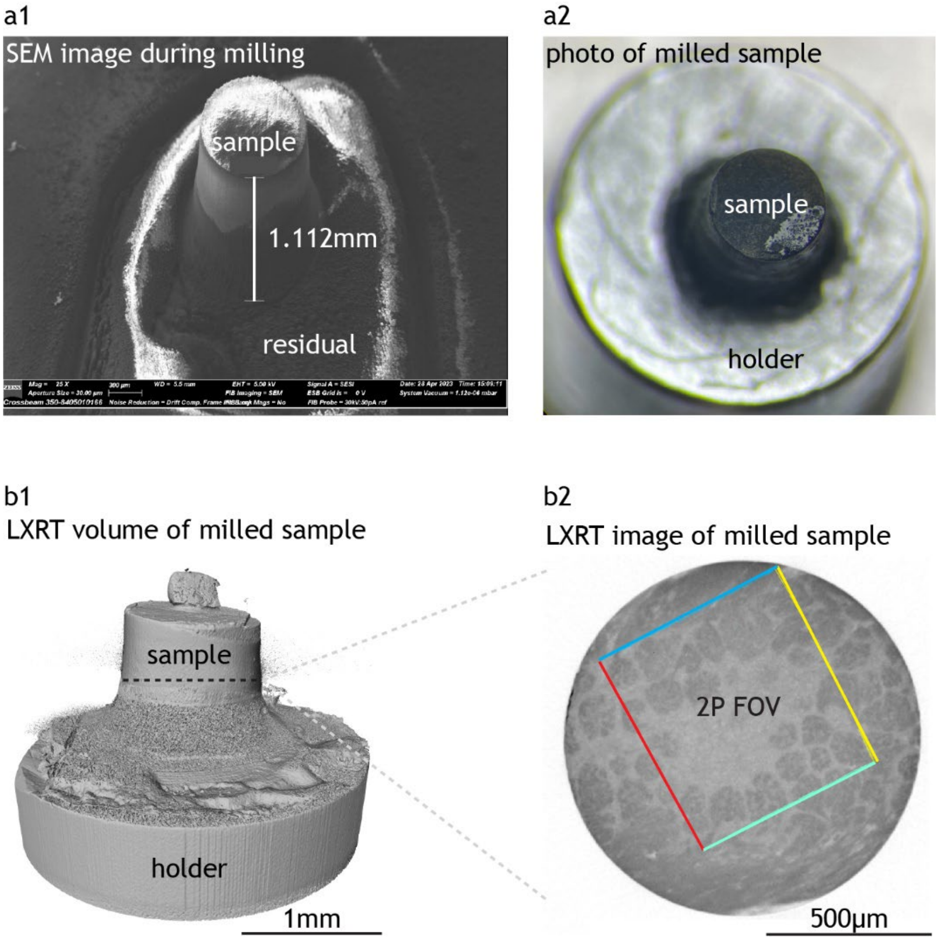

**Supp Fig 1_3: fs-laser milling of sample for XNH.**

**(a)** Scanning electron microscopy (SEM) image **(a1)** and photo **(a2)** of the fs-laser milled cylindrical sample Y489. **(b)** LXRT volume render **(b1)** and example image **(b2)** of the sample, with the location of the 2P-imaged region marked in coloured lines.

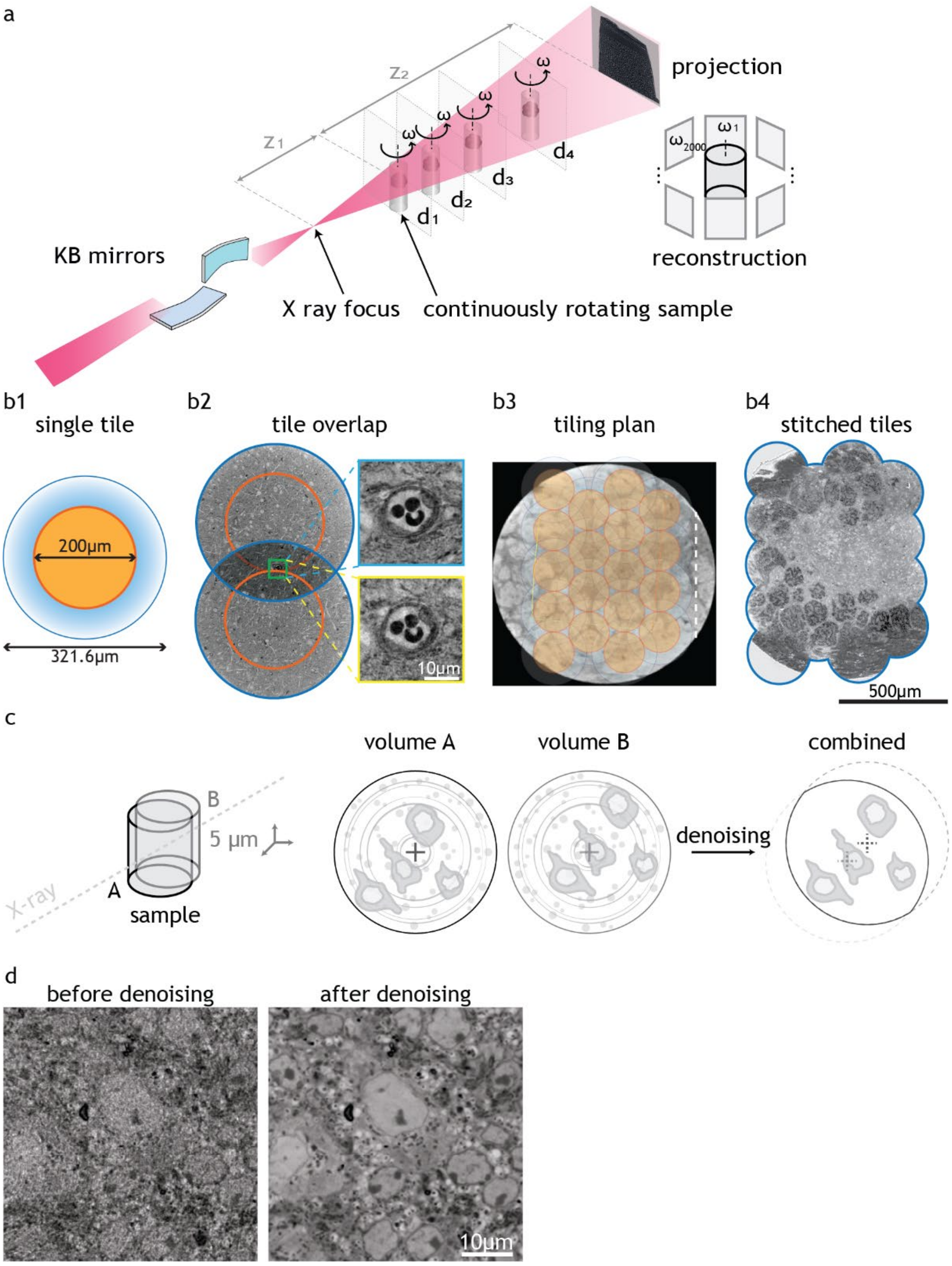

**Supp Fig 1_4: XNH acquisition with continuous acquisition, denoising and tiling.**
**(a)** Imaging principle of XNH (see Methods, Supp Table 3). **(b)** Tiling of XNH field-of-views. **(b1)** Each tile contains a high resolution core (orange, diameter and depth = 200 µm) and an edge that gradually decreases in resolution (blue, diameter and depth = 321.6 µm). **(b2)** Neighbouring tiles were placed with enough overlap to identify common features for subsequent stitching. **(b3)** Tiling plan that covered the 2P-imaged region in correlative sample Y489. **(b4)** Example image after all tiles had been stitched into a continuous volume. **(c)** Denoising principle (see Methods and ref [72]). The sample was acquired twice with a 5 µm shift in stage x, y and z. In the reconstructed volumes, sample features were correlated while imaging noise and ring artefacts were uncorrelated, which allowed them to be removed. **(d)** Image quality comparison before and after dataset denoising (see Methods).

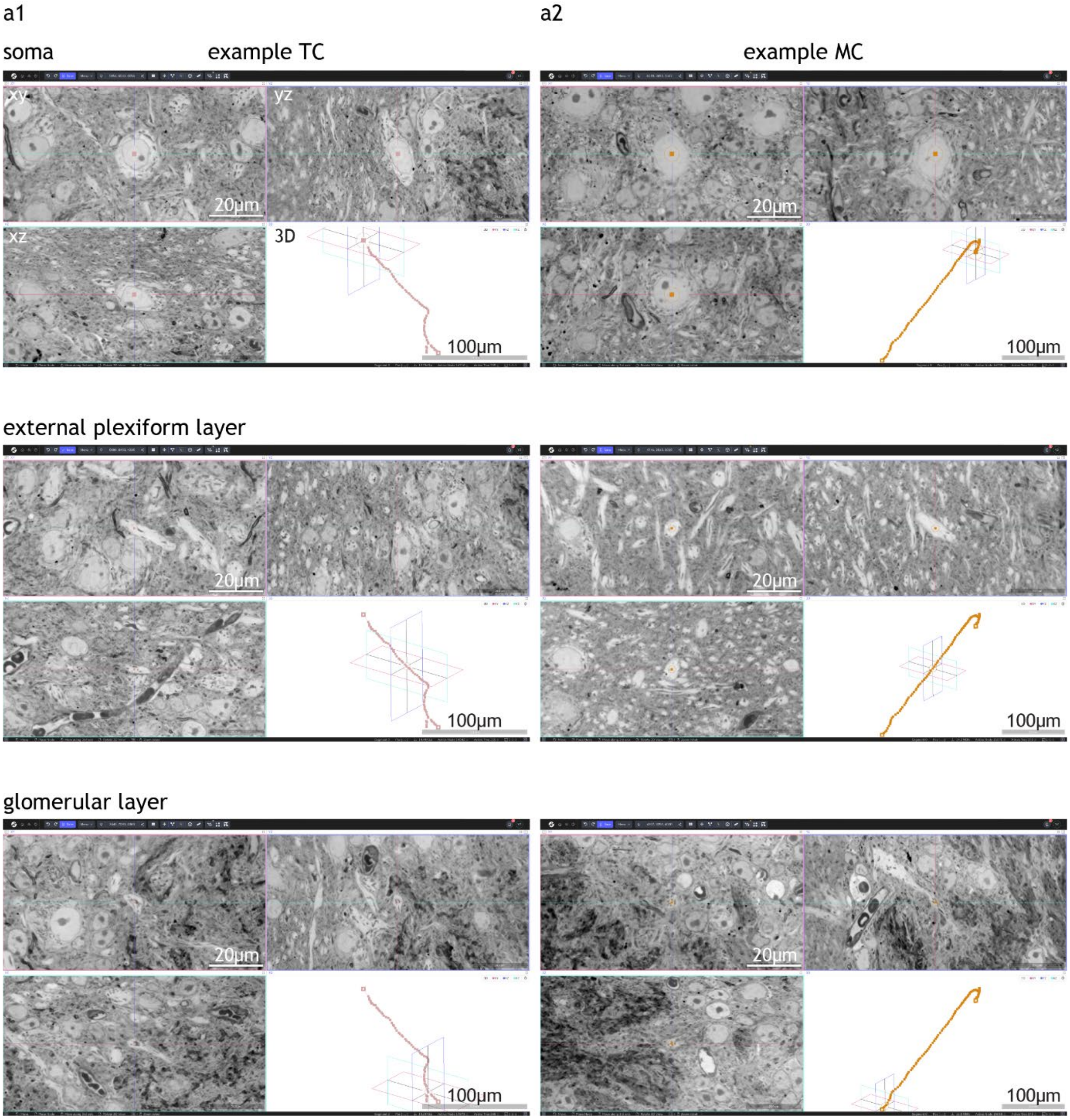

**Supp Fig 1_5: Apical dendrite tracing in XNH datasets.**
**(a)** Screenshots showing the apical dendrite tracing process of an example TC **(a1)** and MC **(a2)** in webKnossos. Each apical dendrite tracing consists of nodes and edges that connect them. Starting from the soma (top), nodes were seeded along the dendrite (middle) until the tracing reached the parent glomerulus of the cell (bottom). See also **Supp Videos 1, 2.**

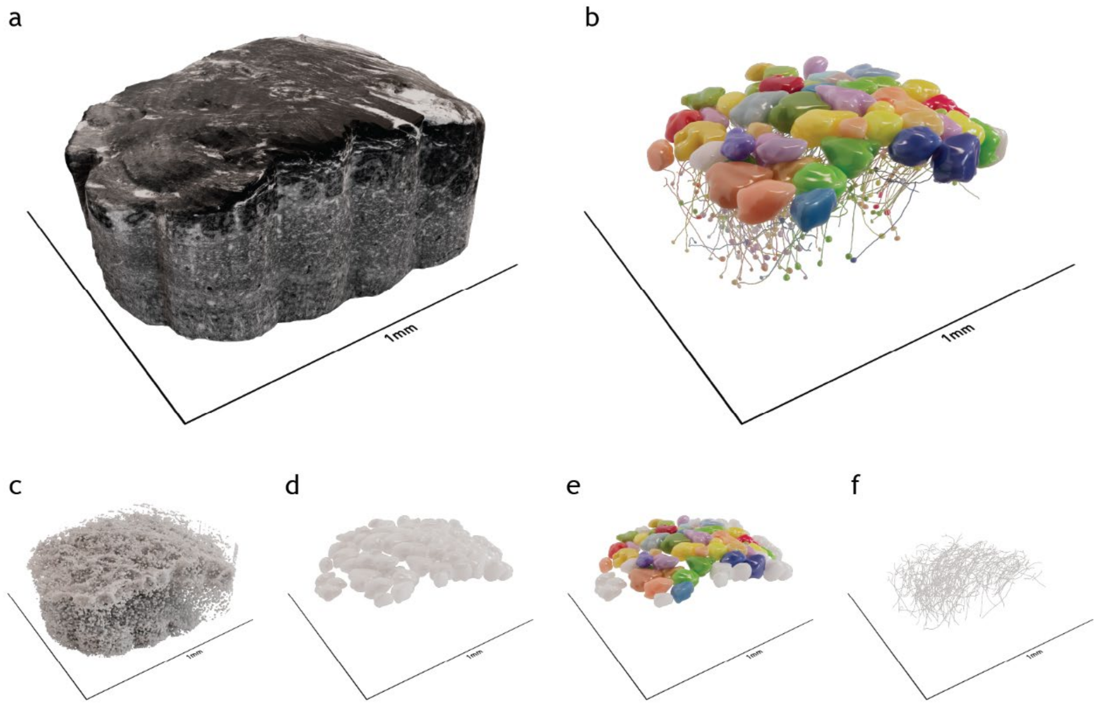

**Supp Fig 1_6: Segmentation of glomeruli, nuclei and apical dendrites of projection neurons in XNH datasets.**

**(a)** 3D render of an example XNH dataset (sample Y489). **(b)** 2P-imaged glomeruli and cells within the XNH volume. Individual glomeruli are uniquely coloured. Each projection neuron (its nucleus and apical dendrite shown) shares the colour of its parent glomerulus. **(c)** Automated segmentation of all cell nuclei. **(d-e)** Consensus manual segmentation of all glomeruli within the XNH volume **(d)**, with those imaged *in vivo* coloured **(e)**. **(f)** Consensus manual skeletonization of the apical dendrites of all projection neurons imaged *in vivo*. See also **Supp Videos 3, 4.**

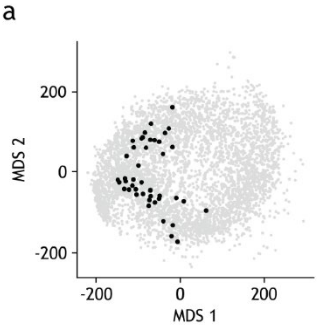

**Supp Fig 1_7: Odorants in chemical space.**
**(a)** Odorants used in this study (black dots) projected in a low dimensional embedding of chemical space, defined by multidimensional scaling (MDS) of chemical distances calculated from 473 descriptors for 5103 chemicals (grey dots).

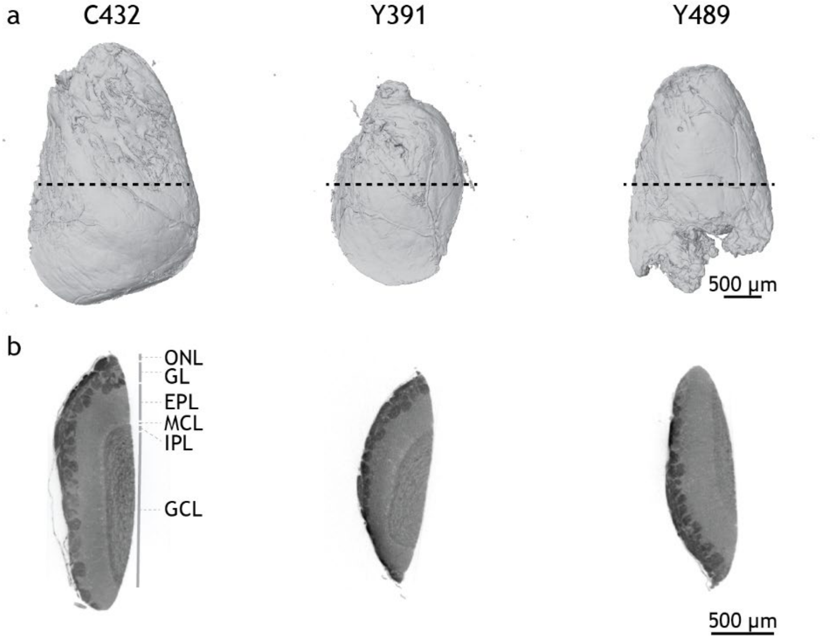

**Supp Fig 1_8: Staining homogeneity assessment using laboratory-based micro-CT (LXRT).**
**(a)** Volume render and example image **(b)** of LXRT datasets of three correlative samples. ONL, olfactory nerve layer; GL, glomerular layer; EPL, external plexiform layer; MCL, mitral cell layer; IPL, internal plexiform layer; GCL, granule cell layer.

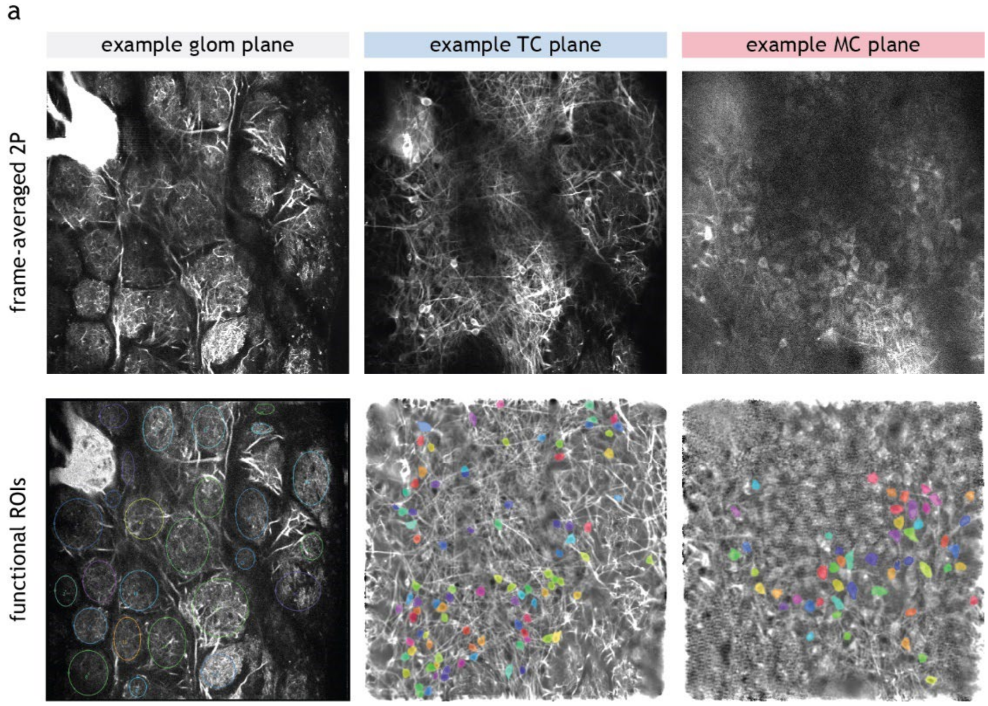

**Supp Fig 1_9: Example 2P images of glomeruli, TC and MC.**
**(a)** Frame-averaged 2P image (n = 880 frames) and functional ROIs of example glomeruli, TC and MC plane from sample Y489.

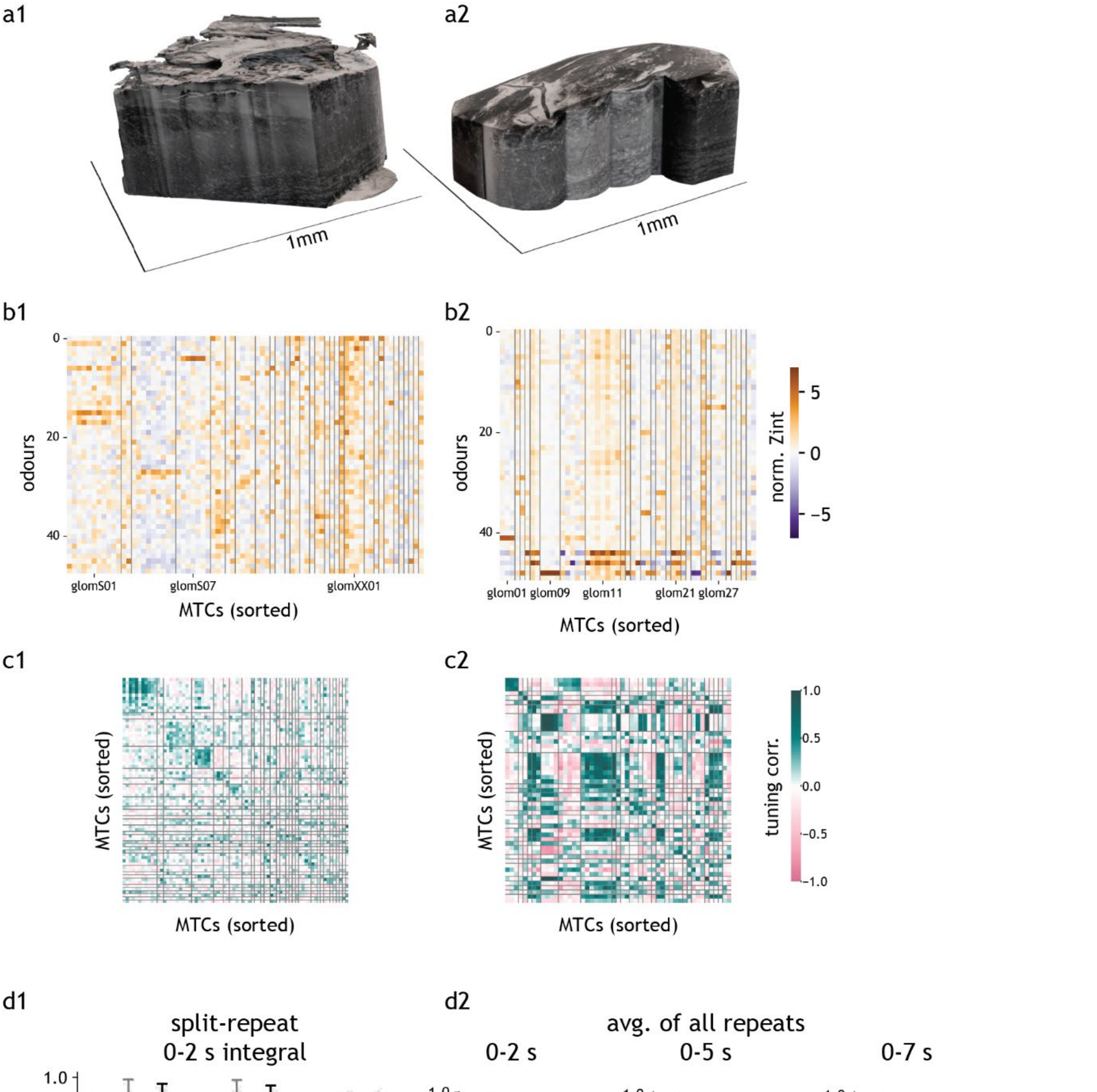

**Supp Fig 2_1: Response matrices of additional correlative datasets.**

**(a)** XNH volume renders of correlative datasets C432 **(a1)** and Y391 **(a2)**. **(b)** Heatmap of odour response integral of cells sorted by parent glomerulus identity for samples C432 **(b1)** and Y391 **(b2)**. Response integral ('Zint') defined as dF/F z-score integral over the odour presentation period (2 s), averaged across all repeats and normalised by the standard deviation of responses to all odours per cell. **(c)** Heatmap of split-repeat odour tuning correlation of cells (see Methods), sorted by parent glomerulus identity for samples C432 **(c1)** and Y391 **(c2)**. **(d)** Tuning correlation summary across three datasets using usual z-score, with split-repeat z-score integral over 2 s integration window **(d1)** or repeat-averaged z-score integral over different integral window sizes for sample Y489 **(d2)**. For the light grey, dark grey and black samples in **d1**, self, sister and non-sister mean values are [0.434, 0.304, 0.034], [0.437, 0.374, 0.002] and [0.522, 0.397, 0.012], respectively and sister > non-sister one-tail t-test *p* values are 0.00029, 4.73E-08 and 2.43E-115, respectively. For 0-2, 0-5 and 0-7 s windows in **d2**, sister and non-sister mean values are [0.550, 0.016], [0.665, 0.061], [0.704, 0.093], respectively and sister > non-sister one-tail t-test *p* values are 6.05E-169, 3.05E-191 and 2.76E-186, respectively. All functionally imaged cells that were traced to their parent glomeruli were used in the analyses of this figure. For **d**, only glomeruli with > 1 sister cell and only strictly responding cells were included (see Methods).

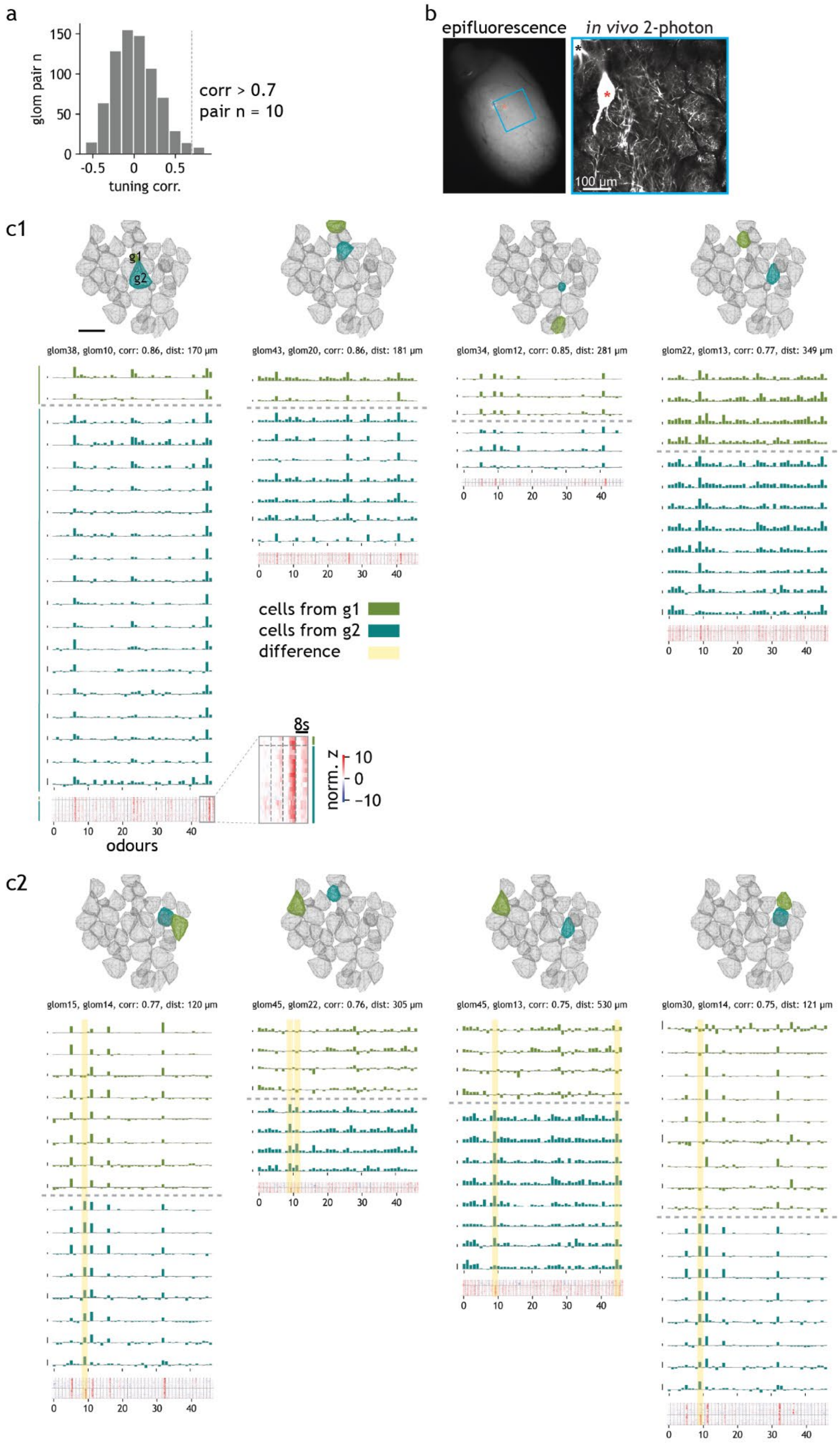

**Supp Fig 2_2: Non-sister cell correlation.**

**(a)** Histogram of averaged sister tuning correlation between glomerulus pairs. Averaged sister tuning refers to the mean odour tuning of all sister cells per glomerulus. **(b)** Epifluorescence and *in vivo* 2P images of a mouse OB with fluorescently labelled MOR174/9 glomerulus split into two (asterisks). **(c)** Bar plot of response integrals (top, scale bar = 10 z-score.) and heatmap of 1 s-binned response z-score (bottom) of sister cells from the top eight glomeruli pairs with the highest tuning correlation. **(c1)** Top correlated glomeruli that have near-identical responses. **(c2)** Top correlated glomerulus pairs that have distinct differences in odour responses (highlighted in yellow). Only glomeruli with > 1 sister cell and only strictly responding cells (see Methods) were included in the analyses of this figure.

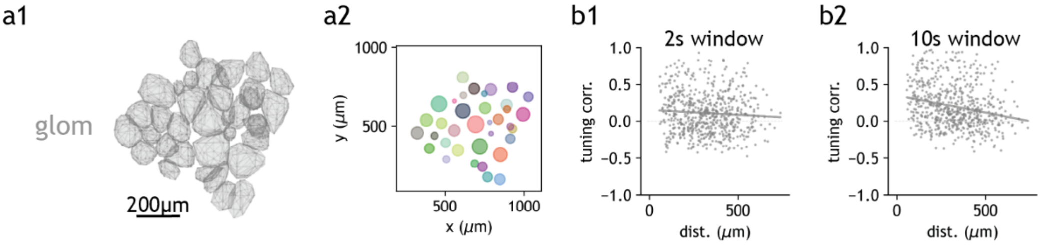

**Supp Fig 2_3: Distance and tuning correlation of glomerulus pairs.**
**(a)** Contours of glomeruli segmented in XNH dataset **(a1)** and represented as dots with diameter proportional to glomerulus volume **(a2)**. **(b)** Odour tuning correlation between glomeruli using 2 s integral (**b1**, $r$ = -0.069, $p$ = 0.083) or 10 s integral (**b2**, $r$ = -0.235, $p$ = 2.42E-9) against centroid-centroid distance between glomerulus pairs. Only glomeruli with 2P data were used in the analyses of this figure.

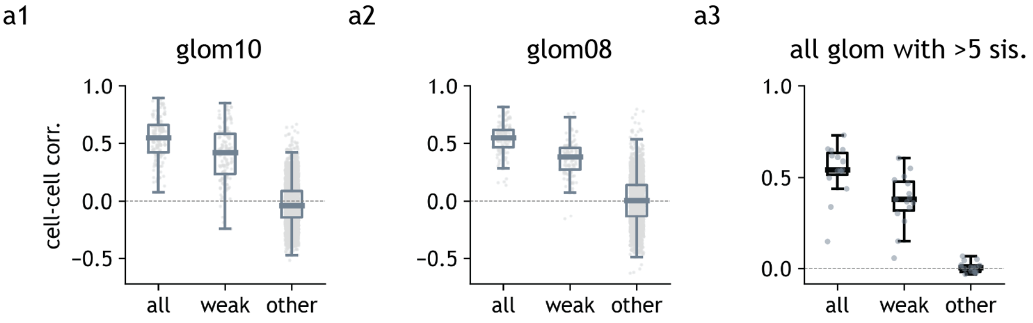

**Supp Fig 3_1: Weak odour correlation in sister cell pairs.**
**(a)** Boxplot of tuning correlation between cell pairs of example glomeruli **(a1-2)** and the mean values pooled across all glomeruli with > 5 sister cells (n = 15) **(a3)**. Three categories represent tuning correlation between sister cell pairs, calculated using all odours ('all') or only weak odours ('weak'); or between non-sister cell pairs using weak odours ('other'). Box = first and third quartile, midline = median, whiskers = most extreme, non-outlier data points. For **a1-3**, the mean values for 'all', 'weak' and 'other' categories are [0.542, 0.41, -0.026], [0.538, 0.367, 0.004] and [0.537, 0.375, 0.007], respectively and the 'weak' > 'other' one-tail t-test $p$ values are 2.08E-156, 1.07E-75 and 1.14E-10, respectively. Only glomeruli with 2P data and > 5 sister cells and only strictly responding cells (see Methods) were used in the analyses of this figure.

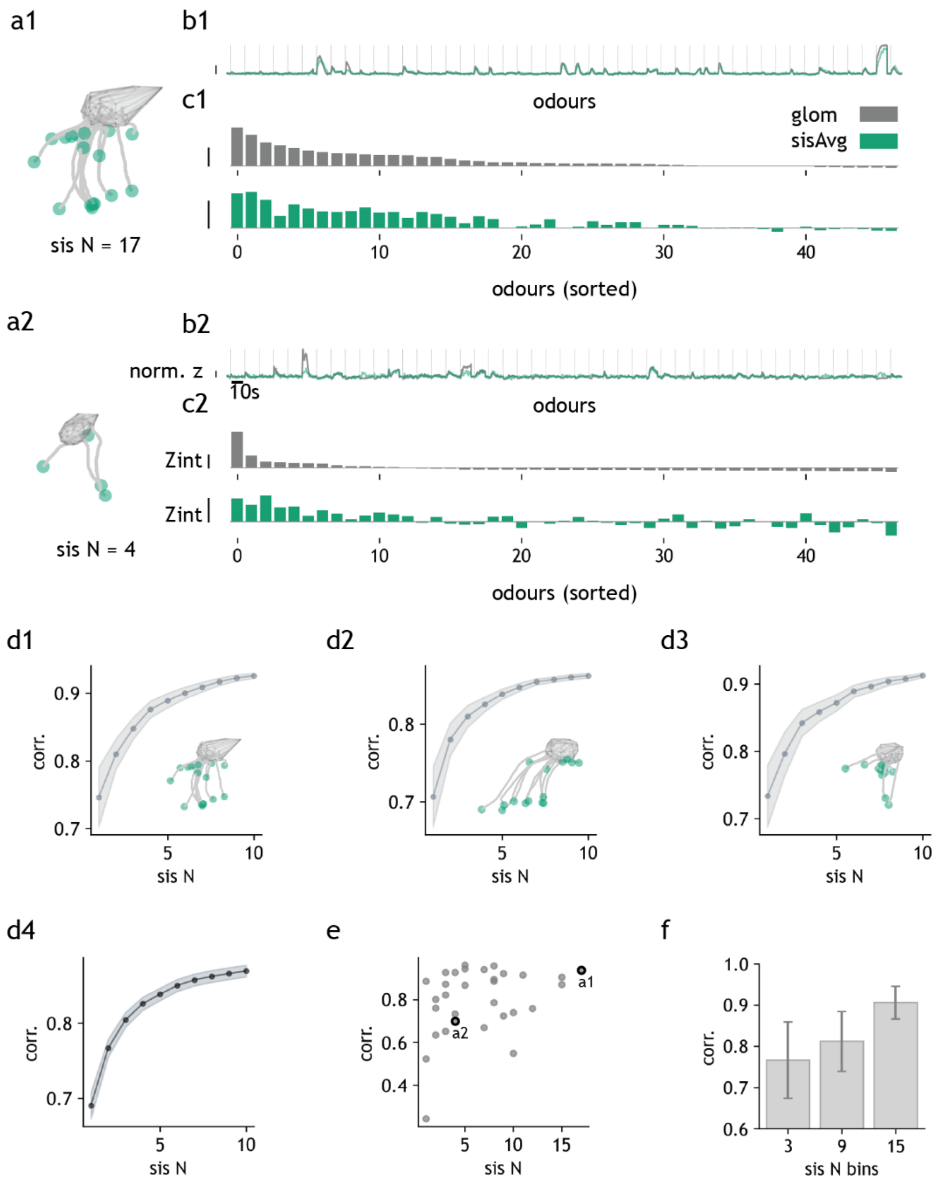

**Supp Fig 3_2: Comparison of glomerular and somatic Tbet-GCaMP6f signal.**
**(a)** Two example glomeruli with high **(a1)** and low **(a2)** number of functionally recorded sister cells. **(b)** Odour response trace (t = -3 to 10s) overlay of glomerulus (grey) and the mean across sister cells (green) for glomeruli in **(a)**. Odours concatenated one after another with onsets marked by vertical lines (t = 0). Scale bars = 2 z-score, normalised by the standard deviation of each glomerulus or cell. **(c)** Bar plot of odour response integrals of parent glomerulus (grey) and the mean across sister cells (green) for glomeruli in **(a)** (scale bars = 2). **(d)** Tuning correlation between parent glomerulus and the mean across sister cells as a function of sister cell number for three example glomeruli **(d1-3)** (shaded area is the 95% confidence interval from 50 random sister cell samplings for each data point) and the mean across glomeruli **(d4)** (shaded areas are the 95% confidence interval of the mean). **(e)** Tuning correlation against total sister number for all glomeruli with 2P data. **(f)** Bar plot of **(e)**, with error bar representing the 95% confidence interval of the mean correlation for each bin. Only glomeruli with 2P data and > 5 sister cells and only strictly responding cells (see Methods) were used in the analyses of this figure.

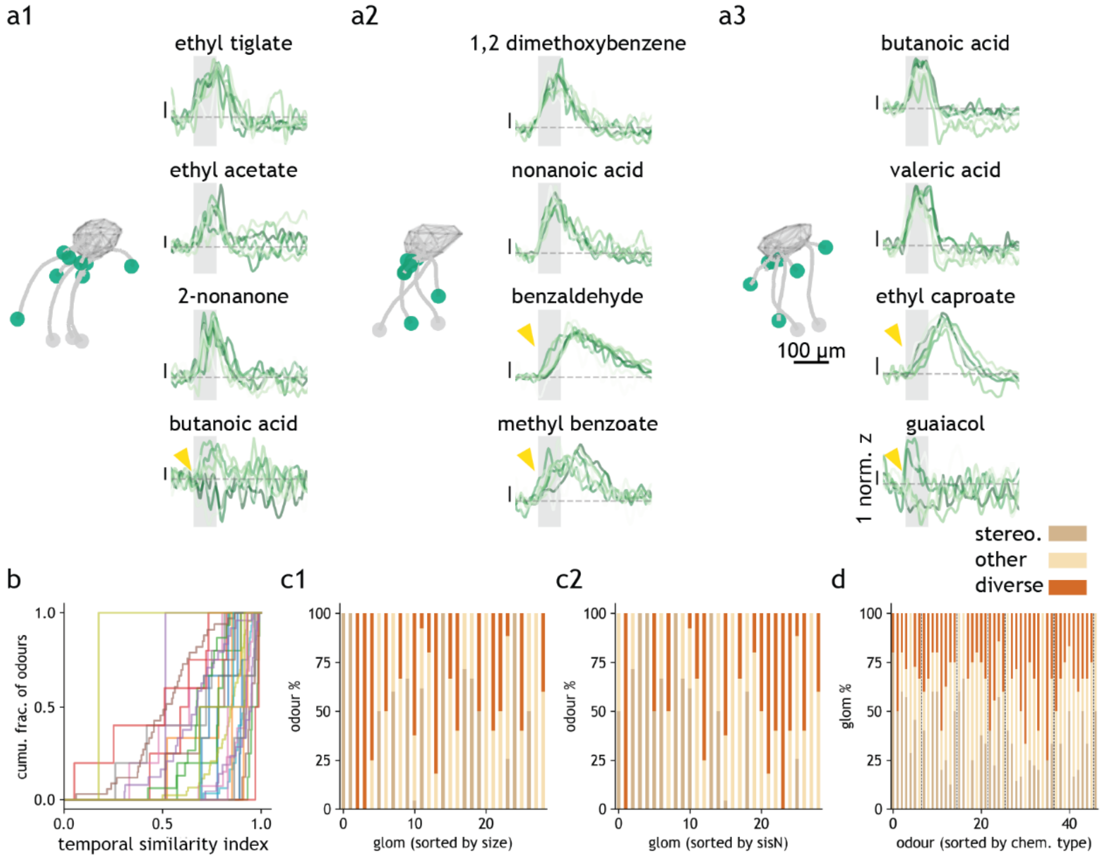

**Supp Fig 4_1: Differences in sister cell responses.**
**(a)** Odour response shape overlay of sister cells (strictly responding [see Methods], green) from three example glomeruli **(a1-3)**. For each cell-odour pair, response shape was defined as repeat-averaged dF/F z-score trace normalised by its standard deviation. Yellow arrowheads mark odours with diverse sister cell responses. **(b)** Cumulative histogram of temporal similarity index (see Methods) of glomerulus-odour pairs for each glomerulus (coloured lines). An index <0.7 is considered diverse and ≥0.9 stereotypical. **(c)** Fraction of stereotypical or diverse odours for each glomerulus, sorted by physical volume **(c1)** or recorded sister cell number **(c2)**. Fraction of stereotypical or diverse glomeruli for each odour, sorted by chemical type. Only glomeruli with multiple sister cells and only strictly responding cells (see Methods) were used in the analyses of this figure. For **(b-c)**, only odours that reliably elicited responses in at least 1 sister cell per glomerulus included.

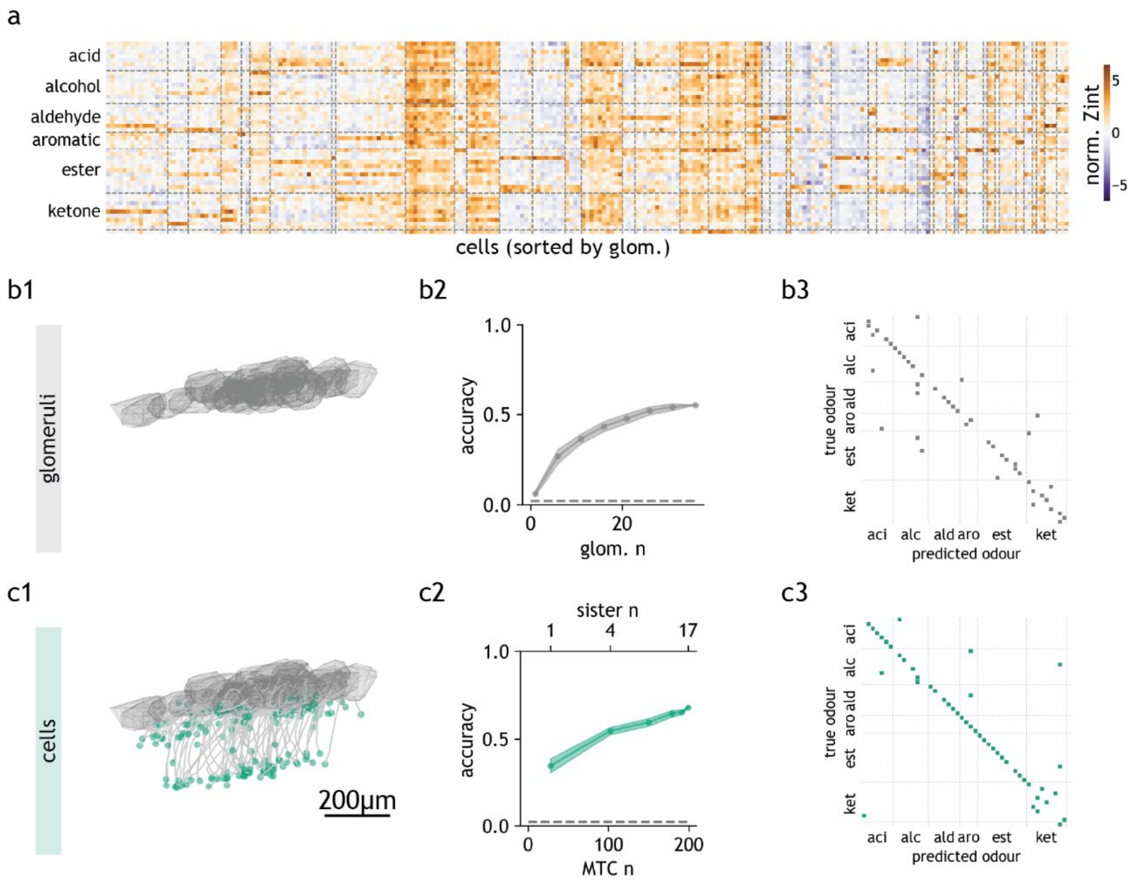

**Supp Fig 4_2: Odour representation and decoding.**

**(a)** Heatmap of odour response integral of cells sorted by parent glomerulus identity and odours sorted by functional group. **(b)** Odour decoding with glomerular responses. **(b1)** Glomeruli contour. **(b2)** Odour decoding accuracy as a function of glomerulus number, from 1 up to 28. The last data point contains all glomeruli in **(b1)**. **(b3)** Confusion matrix of prediction result of the last datapoint in **b2**, with odours sorted by functional group. Prediction accuracies with actual vs shuffled labels are 0.553 and 0.021. **(c)** Odour decoding with somatic responses. **(c1)** Apical dendrite tracing of cells. **(c2)** Odour decoding accuracy as a function of increasing sister cell number, from 1 up to 17 sister cells per glomerulus. The last data point contains all cells in **(c1)**. **(c3)** Confusion matrix of prediction result of the last datapoint in **c2**, with odours sorted by functional group. Prediction accuracies with actual vs shuffled labels are 0.681 and 0.023. **(a)** includes all glomeruli and their strictly responding cells (see Methods), **(b-c)** only includes glomeruli with 2P and > 1 sister cell and strictly responding cells.

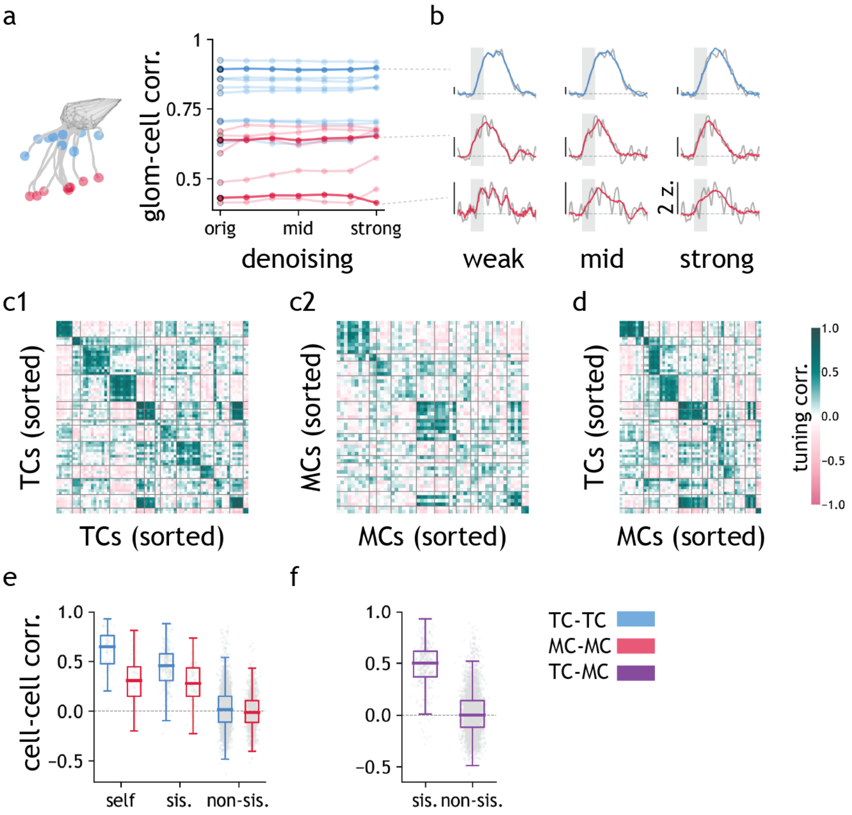

**Supp Fig 5_1: Tuning correlation of sister TCs and sister MCs.**

**(a)** Parent glomerulus-cell tuning correlation as a function of denoising extent (see Methods) for TCs (blue) and MCs (red) of an example glomerulus. **(b)** Overlay of the original (grey) and denoised (coloured) traces for three example cells from **(a)**. **(c)** Odour tuning correlation between TCs (**c1**, split-repeat), MCs (**c2**, split repeat) and between TCs and MCs (**d**, repeat-averaged). **(e)** Boxplot of split-repeat tuning correlation values for each cell across repeats ('self') and for sister and non-sister TC pairs (blue) and MC pairs (red). For TC and MC, self, sister and non-sister mean values are [0.608, 0.441, 0.03] and [0.332, 0.277, -0.001], respectively and sister > non-sister one-tail t-test *p* values are 7.99E-158 and 3.80E-54, respectively. **(f)** Boxplot of repeat-averaged tuning correlation values between sister and non-sister TC-MC pairs. (box = first and third quartile, midline = median, whiskers = most extreme, non-outlier data points. Sister and non-sister mean values are 0.48 and 0.012; sister >non-sister one tail t-test *p* value is 2.38E-275.) Only glomeruli with 2P data and > 1 TC and > 1 MC and only strictly responding cells (see Methods) were used in the analyses of this figure.

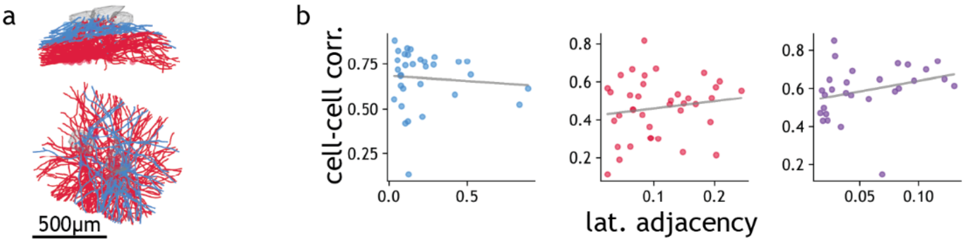

**Supp Fig 5_2: Tuning correlation and lateral dendrite proximity.**
**(a)** Side and dorsal view of lateral dendrite tracings of three example glomeruli. **(b)** Relationship between cell-cell tuning correlation and lateral adjacency score for TC-TC ($r$ = -0.078, $p$ = 0.669), MC-MC ($r$ = 0.149, $p$ = 0.372) and TC-MC ($r$ = 0.286, $p$ = 0.140) cell pairs, pooled across glomeruli in **(a)**. Only includes three glomeruli with fully traced lateral dendrites. Only including strictly responding cells and cell pairs with non-zero lateral adjacency score (see Methods).

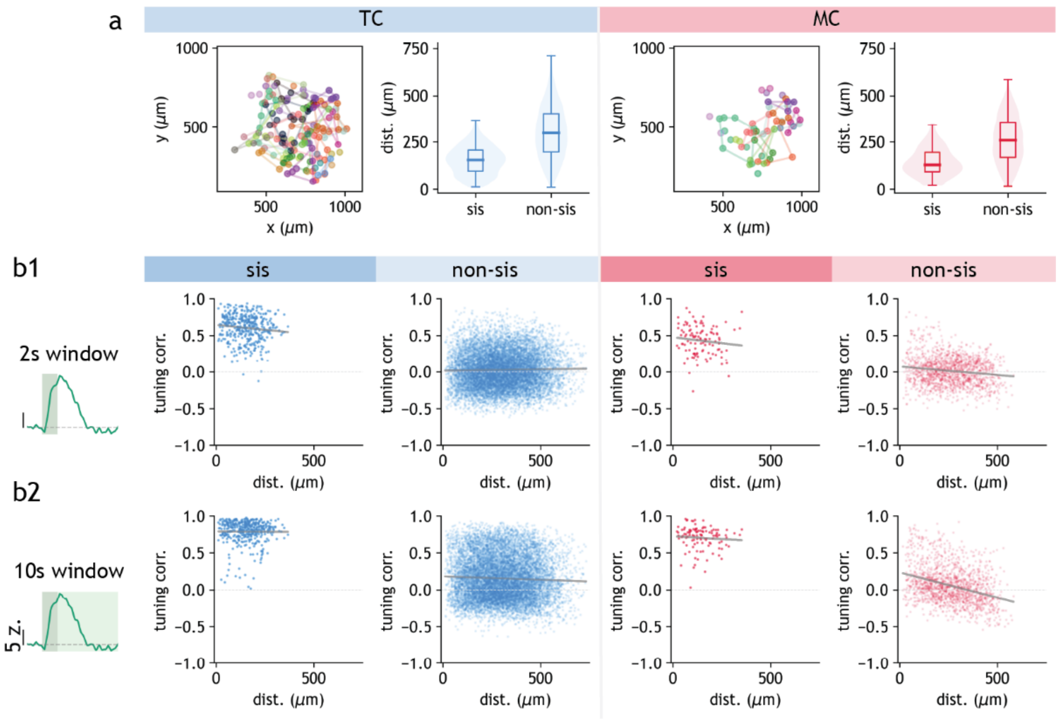

**Supp Fig 5_3: Distance and tuning correlation of TC pairs and MC pairs.**
**(a)** Location of TCs and MCs, coloured and enclosed by parent glomerulus identity as in Supp Fig F2_3, and boxplot of somata distances between sister and non-sister TCs and MCs (box = first and third quartile, midline = median, whiskers = most extreme, non-outlier data points). **(b)** Odour tuning correlation against soma distances of sister and non-sister TCs and MCs using 2 s **(b1)** or 10 s **(b2)** response integral. The correlation coefficients and $p$ values are as follows. 'TC, sis, 2s': $r$ = -0.108, $p$ = 0.031; 'TC, sis, 10s': $r$ = -0.007, $p$ = 0.885; 'TC, non-sis, 2s': $r$ = 0.019, $p$ = 0.053; 'TC, non-sis, 10s': $r$ = -0.044, $p$ = 4.50E-06; MC, sis, 2s': $r$ = -0.115, $p$ = 0.235; 'MC, sis, 10s': $r$ = -0.070, $p$ = 0.470; 'MC, non-sis, 2s': $r$ = -0.147, $p$ = 1.22E-08; 'MC, non-sis, 10s': $r$ = -0.340, $p$ = 1.94E-41. Only glomeruli with 2P data and > 1 TC and > 1 MC and only strictly responding cells (see Methods) were used in the analyses of this figure.

a

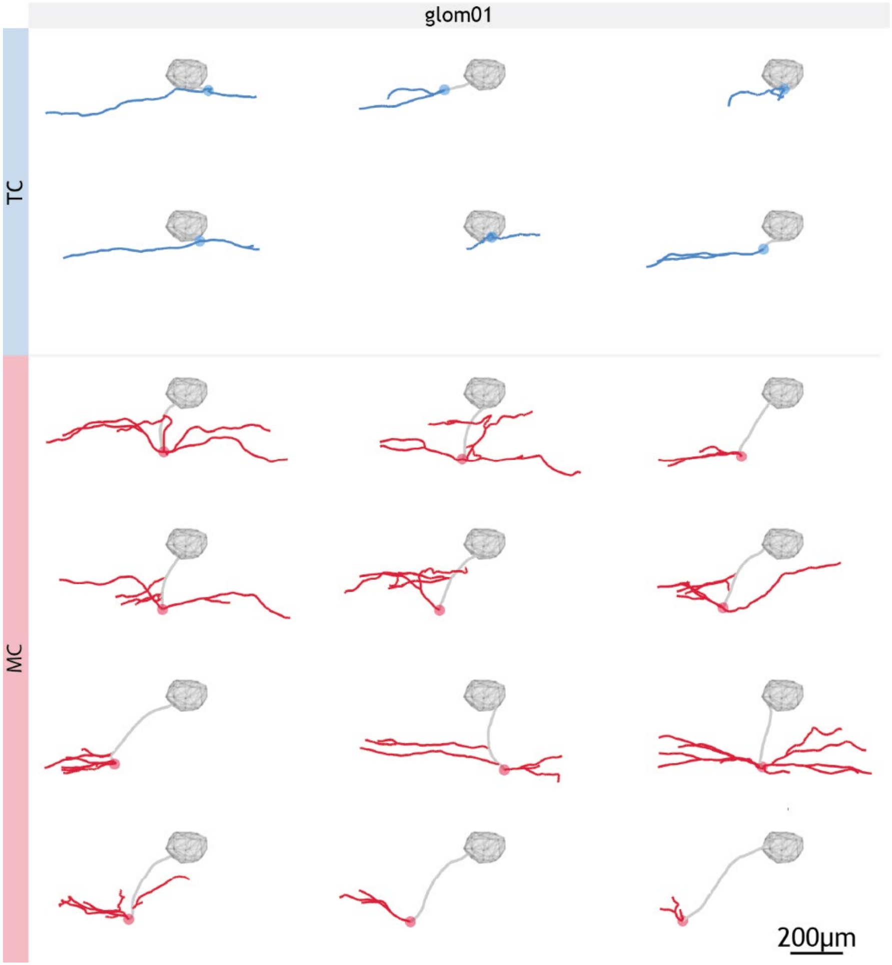

**Supp Fig 5_4: Gallery of example apical and lateral dendrite tracings 1.**
**(a)** Apical dendrite (grey line) and lateral dendrite (coloured line) tracings of sister TCs (blue) and MCs (red) to their parent glomerulus ‘glom01’.

a

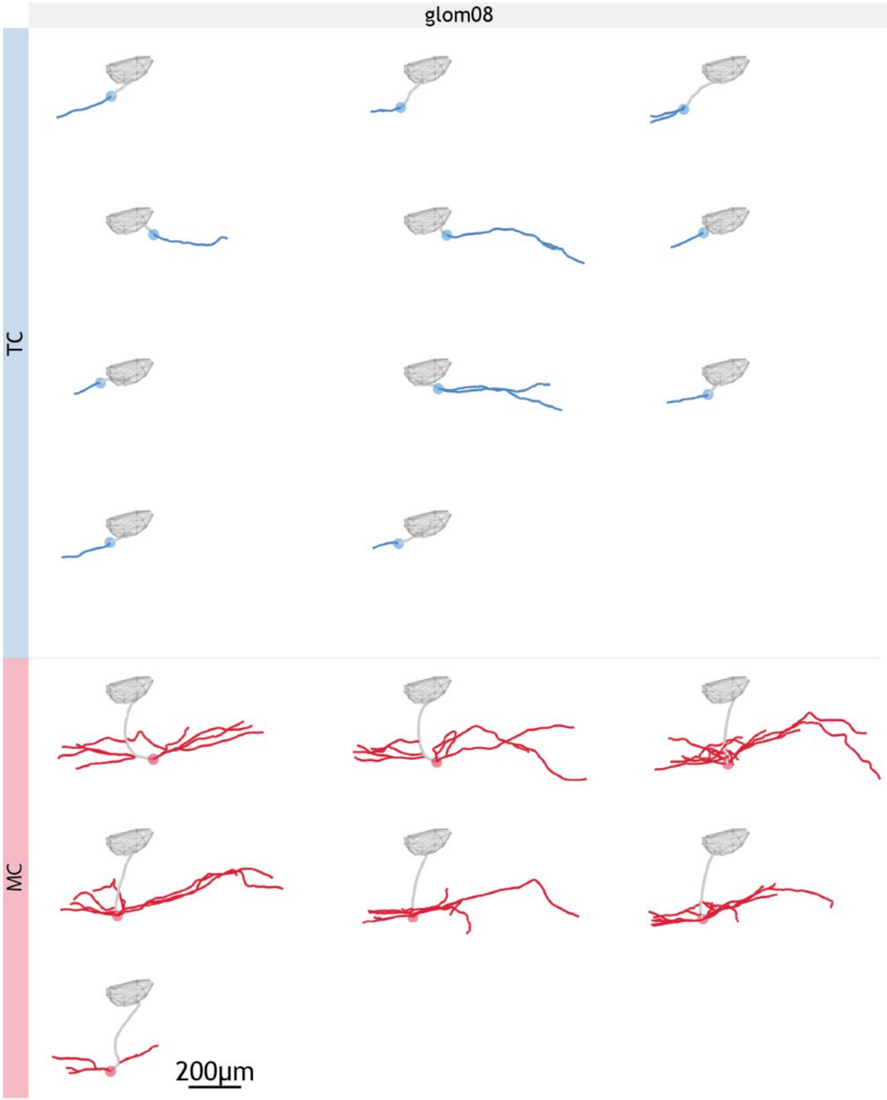

**Supp Fig 5_5: Gallery of example apical and lateral dendrite tracings 2.**
**(a)** Apical dendrite (grey line) and lateral dendrite (coloured line) tracings of sister TCs (blue) and MCs (red) to their parent glomerulus ‘glom08’.

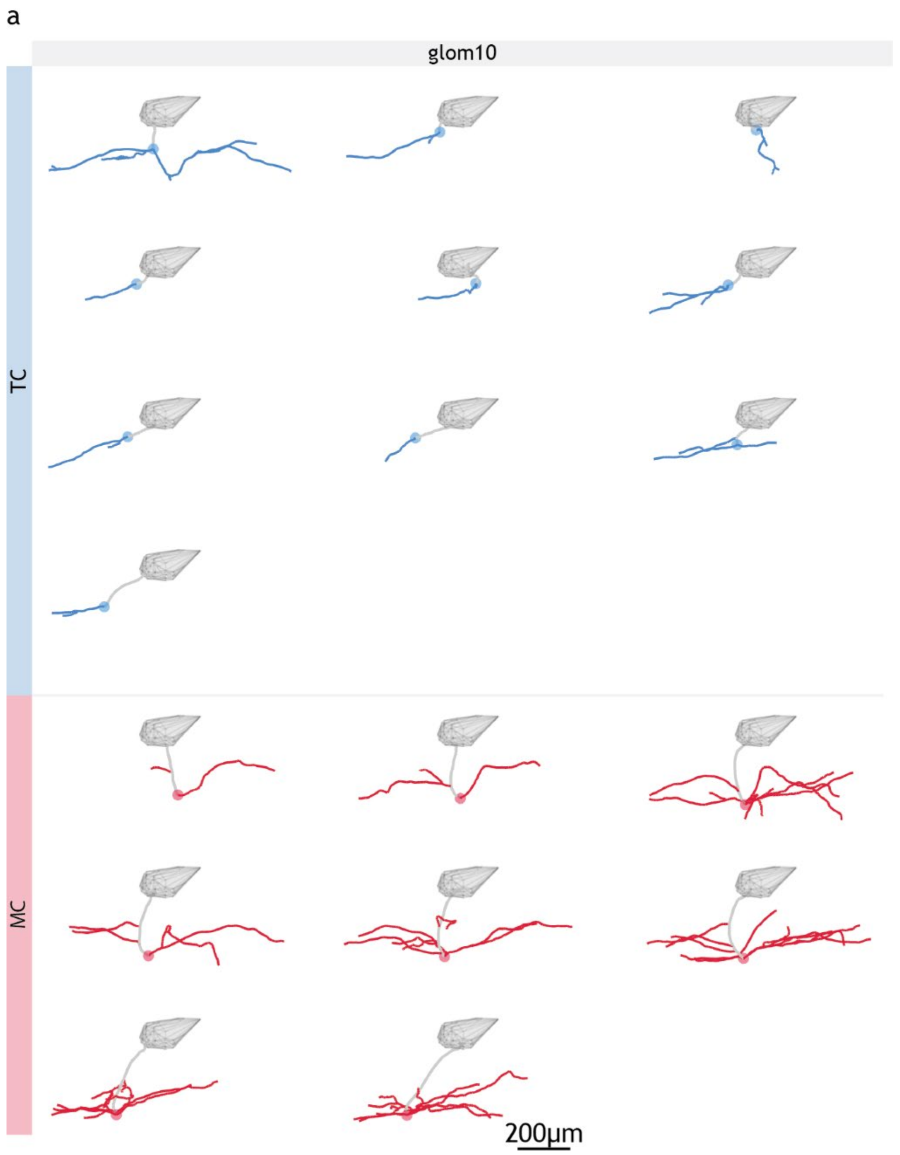

**Supp Fig 5_6: Gallery of example apical and lateral dendrite tracings 3.**
**(a)** Apical dendrite (grey line) and lateral dendrite (coloured line) tracings of sister TCs (blue) and MCs (red) to their parent glomerulus ‘glom10’. One MC had no traceable lateral dendrites and is not included.

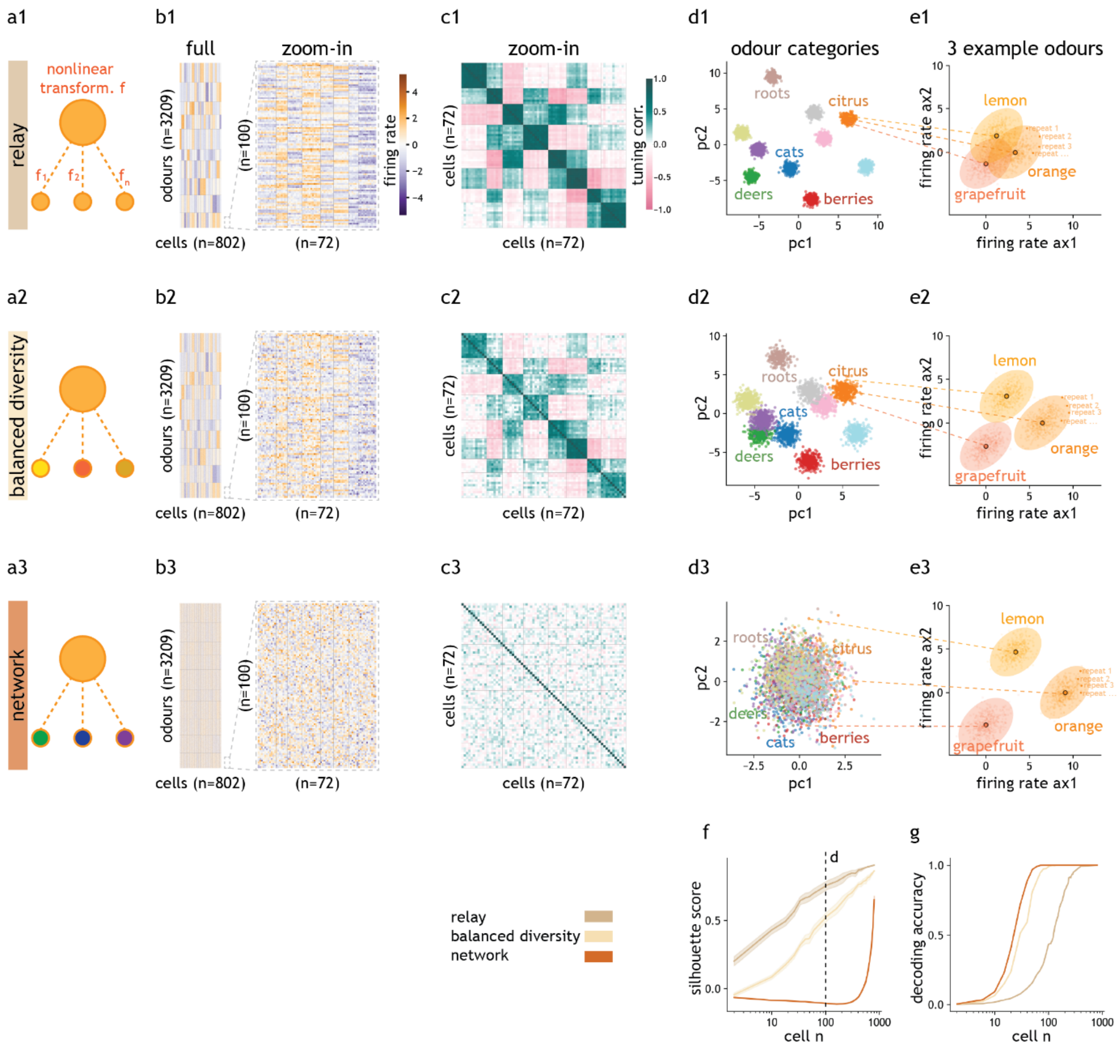

**Supp Fig 5_7: Odour encoding and generalization capacity of nonlinear sister cell models.**

**(a)** Sister cell models, where sister cells are increasingly dissimilar to each other and to the parent glomerulus from model relay **(a1)**, balanced diversity **(a2)** to network **(a3)**. Cell activity modelled as nonlinear transformations of glomerulus activity (see Methods). **(b)** Simulated response matrix of 802 sister cells, sorted by parent glomerulus (n = 20) to 3209 odours, sorted by categories (n = 10), and the zoom-in view of a subset of the matrix, for each model. **(c)** Odour tuning correlation of cells from the zoomed in view in **(b)** showing high, medium and low sister correlation in models relay **(c1)**, balanced diversity **(c2)** and network **(c3)**, respectively. **(d)** Odour generalization ability visualized by the projection of response vectors of all odours to the top two principal components of cell space, where both model relay **(d1)** balanced diversity **(d2)** show clustering of odour categories in contrast to model network which does not **(d3)**. **(e)** Odour encoding capacity visualized by the degree of overlap in the ‘noise cloud’ (i.e. 500 repeats) of three odours from the same category, where both model balanced diversity **(e2)** and network **(e3)** shows clear noise cloud separation in contrast to model relay which does not **(e1)**. **(f)** Quantification of odour category clustering in **(d)** by average silhouette score, as a function of cell number. 100 cells were used in the PCA projection in **(d)**, as marked by the dashed line. Shading represents 95% confidence interval calculated from 20 independent cell samples. **(g)** Quantification of odour encoding capacity as exampled in **(e)** by odour decoding accuracy with linear SVM, as a function of cell number.

### Appendix:

**Appendix 1 Details of the toy model.**

In the sister cell toy model (**Supp Fig 5_7**), projection neuron activity was modelled as a non-linear transformation of the input they receive, which was in turn modelled as a noisy version of the parent glomerulus activity. The singular value decomposition of cell input activity matrix $Y = USV^T$ and cell output matrix $X = URV^T$ satisfies $R = \frac{S \pm \sqrt{S^2 + 8/\beta}}{2}$. The only free parameter β controls the degree of the sister cell similarity. When β is large, sister cells have similar and hence correlated odour responses and when β is small, sister cells have very uncorrelated odour responses (see Methods).

The above input-output transformation is the closed-form solution of a loss function minimization problem aimed at maximizing both the odour encoding capacity and novel odour generalization ability of the projection neuron population, which will be explained below.

In nature, animals encounter a huge variety of different odours. Some odours are more perceptually similar than others and this probably reflects an ethological need or relevance to generalise across certain smells (e.g. leaves, rotting fruits). This perceptual odour landscape may also help the animal to infer the property of novel objects based on its smell (e.g. a new kind of citrus fruits). An ideal olfactory system should, among other things, be able to represent a large number of odours in a way that they can be easily discriminated, while retaining a meaningful structure in this odour representation to allow generalization of novel odours to potentially related odour concepts. In this toy model, the odour encoding capacity was approximated as the representational volume of the cell output matrix *X*. If we plot each odour as a dot in the high dimensional space defined by the firing rate of cells, the larger the volume occupied by odours, the more spread out the odour representations will tend to be, and the higher the encoding capacity of the cells, as measured by the number of possible discriminations. Hence, to maximize encoding capacity, we want to maximize $vol(X) = \prod(\sigma_i) = \sqrt{det(XX^T)}$. Odour generalization ability was in turn approximated by how much the cell population can retain the chemical similarity of odours. Chemical similarity was assumed to be equivalent to glomerular response similarity between odours. The system will be able to generalize if it can correctly assign a novel odour to its closest odour category.

To mimic odour categories, odours with similar glomerular responses were simulated. In glomerulus representation space, odours are clustered. If such structure is preserved in cell representation space, we would consider the system to have high generalization ability, whereby a novel odour will be represented in the cluster of the highest chemical similarity. One way to achieve this, and the way adopted by this toy model, is to enforce the cell output *X* to be similar to the glomeruli activity (equivalent to cell input *Y*) i.e. minimizing $||X - Y||^2$.

The two terms can be combined into a loss function $L(X) = -log(det(XX^T)) + 0.5\beta||X - Y||^2$ where β controls the relative importance of maximizing odour encoding capacity versus preserving chemical similarity (i.e. odour generalization ability). An output representation *X* can be found by minimizing this loss function with respect to *X*. The gradient of this loss function is $\nabla_x L = -2(XX^T)^{-1}X + \beta(X - Y)$. Setting $\nabla_x L = 0$, we have:

$$
\begin{aligned}
2(XX^T)^{-1}X &= \beta(X-Y)\\
2X &= \beta(XX^T)(X-Y)\\
2X &= \beta(XX^T)X - \beta(XX^T)Y\\
2XX^T &= \beta(XX^T)XX^T - \beta(XX^T)YX^T\\
&\text{multiply by } (XX^T)^{-1}\text{on each side}\\
2I &= \beta(XX^T) - \beta(YX^T)\\
\beta YX^T &= \beta XX^T - 2I
\end{aligned}
$$

One solution to this equation occurs when *X* and *Y* share the same rotational matrices *U* and *V*.

Hence, we can substitute *X* and *Y* with:

$$X = URV^T$$
$$Y = USV^T$$

Which will give:

$$
\begin{aligned}
\beta US(V^TV)RU^T &= \beta UR(V^TV)RU^T - 2I\\
\beta USRU^T &= \beta URRU^T - 2I\\
\beta(U^TU)SRU^T &= \beta(U^TU)RRU^T - 2U^T\\
\beta SRU^T &= \beta RRU^T - 2U^T\\
\beta SR(U^TU) &= \beta RR(U^TU) - 2(U^TU)\\
\beta SR &= \beta RR - 2\\
0 &= R^2 - SR - 2/\beta
\end{aligned}
$$

The root of this quadratic equation $R = \frac{S \pm \sqrt{S^2 + 8/\beta}}{2}$ defines the non-linear input-output transformation of the projection neuron population.

### Supplementary tables:

*sample C432*

| idx | moPairs | odourant |
|---|---|---|
| 0 | modd01_odour1 | Nonanoic Acid |
| 1 | modd01_odour2 | 2-Hydroxyacetophenone |
| 2 | modd01_odour3 | 1-Nonanol |
| 3 | modd01_odour4 | 1-Heptanol |
| 4 | modd01_odour5 | 1,2-Dimethoxybenzene |
| 5 | modd01_odour6 | (+)-Fenchone |
| 6 | modd02_odour1 | 2-Phenylpropionaldehyde |
| 7 | modd02_odour2 | 2-Nonanone |
| 8 | modd02_odour3 | 2-Methyl-2-Butanol |
| 9 | modd02_odour4 | 2-Methoxy-4-Methylphenol |
| 10 | modd02_odour5 | Hexanoic Acid |
| 11 | modd02_odour6 | 2-Heptanone |
| 12 | modd03_odour1 | Benzyl Acetate |
| 13 | modd03_odour2 | Benzaldehyde |
| 14 | modd03_odour3 | alpha-Terpinene |
| 15 | modd03_odour4 | Acetophenone |
| 16 | modd03_odour5 | 4-Phenyl-2-Butanone |
| 17 | modd03_odour6 | 4-Allylanisole |
| 18 | modd04_odour1 | Ethyl Valerate |
| 19 | modd04_odour2 | Ethyl Heptanoate |
| 20 | modd04_odour3 | Ethyl Butyrate |
| 21 | modd04_odour4 | Ethyl Acetate |
| 22 | modd04_odour5 | Cyclohexanol |
| 23 | modd04_odour6 | cis-3-Hexenyl Tiglate |
| 24 | modd05_odour1 | Propyl Acetate |
| 25 | modd05_odour2 | Isoamyl Acetate |
| 26 | modd05_odour3 | 1,4-Cineole |
| 27 | modd05_odour4 | Guaiacol |
| 28 | modd05_odour5 | Eugenol |
| 29 | modd05_odour6 | Eucalyptol |
| 30 | modd06_odour1 | Methyl Valerate |
| 31 | modd06_odour2 | Methyl Salicylate |
| 32 | modd06_odour3 | Methyl Butyrate |
| 33 | modd06_odour4 | Methyl Benzoate |
| 34 | modd06_odour5 | Geranyl Acetate |
| 35 | modd06_odour6 | Isobutyric Acid |
| 36 | modd07_odour1 | Ethyl Caproate |
| 37 | modd07_odour2 | Valeraldehyde |
| 38 | modd07_odour3 | S-(+)-Carvone |
| 39 | modd07_odour4 | R-(+)-Limonene |
| 40 | modd07_odour5 | Octanal |
| 41 | modd07_odour6 | Valeric Acid |
| 42 | modd08_odour1 | Mineral Oil |
| 43 | modd08_odour2 | 2,4-Dimethylacetophenone |
| 44 | modd08_odour3 | Menthone |
| 45 | modd08_odour4 | Ethyl Tiglate |
| 46 | modd08_odour5 | 2-Hexanone |
| 47 | modd08_odour6 | 4-Methylacetophenone |

*sample Y391*

| idx | moPairs | odourant |
|---|---|---|
| 0 | modd01_odour1 | Nonanoic Acid |
| 1 | modd01_odour2 | 2-Hydroxyacetophenone |
| 2 | modd01_odour3 | 1-Nonanol |
| 3 | modd01_odour4 | 2-Phenylpropionaldehyde |
| 4 | modd01_odour5 | 1,2-Dimethoxybenzene |
| 5 | modd01_odour6 | Ethyl Valerate |
| 6 | modd02_odour1 | Trans-Anethole |
| 7 | modd02_odour2 | 2-Nonanone |
| 8 | modd02_odour3 | 2-Methyl-2-Butanol |
| 9 | modd02_odour4 | Benzaldehyde |
| 10 | modd02_odour5 | Hexanoic Acid |
| 11 | modd02_odour6 | Methyl Valerate |
| 12 | modd03_odour1 | Benzyl Acetate |
| 13 | modd03_odour2 | 1-Heptanol |
| 14 | modd03_odour3 | alpha-Terpinene |
| 15 | modd03_odour4 | Acetophenone |
| 16 | modd03_odour5 | Valeraldehyde |
| 17 | modd03_odour6 | Geranyl Acetate |
| 18 | modd04_odour1 | (+)-Fenchone |
| 19 | modd04_odour2 | Ethyl Heptanoate |
| 20 | modd04_odour3 | 4-Allylanisole |
| 21 | modd04_odour5 | Cyclohexanol |
| 22 | modd04_odour6 | Dodecanal |
| 23 | modd05_odour1 | Propyl Acetate |
| 24 | modd05_odour3 | 1,4-Cineole |
| 25 | modd05_odour4 | Guaiacol |
| 26 | modd06_odour1 | Butanoic Acid |
| 27 | modd06_odour2 | Methyl Salicylate |
| 28 | modd06_odour3 | 2-Methoxy-4-Methylphenol |
| 29 | modd06_odour4 | 2-Heptanone |
| 30 | modd06_odour5 | Nonanal |
| 31 | modd06_odour6 | cis-3-Hexenyl Tiglate |
| 32 | modd07_odour1 | Ethyl Caproate |
| 33 | modd07_odour2 | Eugenol |
| 34 | modd07_odour3 | S-(+)-Carvone |
| 35 | modd07_odour4 | Methyl Benzoate |
| 36 | modd07_odour5 | Octanal |
| 37 | modd07_odour6 | Valeric Acid |
| 38 | modd08_odour1 | Mineral Oil |
| 39 | modd08_odour2 | 2,4-Dimethylacetophenone |
| 40 | modd08_odour3 | Eucalyptol |
| 41 | modd08_odour4 | Ethyl Tiglate |
| 42 | modd08_odour5 | Undecanal |
| 43 | modd08_odour6 | R-Citronellic Acid |
| 44 | modd14_odour1 | Methyl Butyrate |
| 45 | modd14_odour4 | R-(+)-Limonene |
| 46 | modd14_odour6 | Ethyl Butyrate |
| 47 | modd15_odour1 | 4-Methyloctanoic Acid |
| 48 | modd15_odour2 | Ethyl Acetate |
| 49 | modd15_odour3 | 4-Methylacetophenone |

*sampel Y489*

| idx | moPairs | odourant |
|---|---|---|
| 0 | modd01_odour1 | Nonanoic Acid |
| 1 | modd01_odour2 | 2-Hydroxyacetophenone |
| 2 | modd01_odour3 | 1-Nonanol |
| 3 | modd01_odour4 | 2-Phenylpropionaldehyde |
| 4 | modd01_odour5 | 1,2-Dimethoxybenzene |
| 5 | modd01_odour6 | Ethyl Valerate |
| 6 | modd02_odour1 | Trans-Anethole |
| 7 | modd02_odour2 | 2-Nonanone |
| 8 | modd02_odour3 | 2-Methyl-2-Butanol |
| 9 | modd02_odour4 | Benzaldehyde |
| 10 | modd02_odour5 | Hexanoic Acid |
| 11 | modd02_odour6 | Methyl Valerate |
| 12 | modd03_odour1 | Benzyl Acetate |
| 13 | modd03_odour2 | 1-Heptanol |
| 14 | modd03_odour3 | alpha-Terpinene |
| 15 | modd03_odour4 | Acetophenone |
| 16 | modd03_odour5 | Valeraldehyde |
| 17 | modd03_odour6 | Geranyl Acetate |
| 18 | modd04_odour1 | (+)-Fenchone |
| 19 | modd04_odour2 | Ethyl Heptanoate |
| 20 | modd04_odour3 | 4-Allylanisole |
| 21 | modd04_odour5 | Cyclohexanol |
| 22 | modd04_odour6 | Dodecanal |
| 23 | modd05_odour1 | Propyl Acetate |
| 24 | modd05_odour3 | 1,4-Cineole |
| 25 | modd05_odour4 | Guaiacol |
| 26 | modd06_odour1 | Butanoic Acid |
| 27 | modd06_odour2 | Methyl Salicylate |
| 28 | modd06_odour3 | 2-Methoxy-4-Methylphenol |
| 29 | modd06_odour4 | 2-Heptanone |
| 30 | modd06_odour5 | Nonanal |
| 31 | modd06_odour6 | cis-3-Hexenyl Tiglate |
| 32 | modd07_odour1 | Ethyl Caproate |
| 33 | modd07_odour2 | Eugenol |
| 34 | modd07_odour3 | S-(+)-Carvone |
| 35 | modd07_odour4 | Methyl Benzoate |
| 36 | modd07_odour5 | Octanal |
| 37 | modd07_odour6 | Valeric Acid |
| 38 | modd08_odour1 | Mineral Oil |
| 39 | modd08_odour2 | 2,4-Dimethylacetophenone |
| 40 | modd08_odour3 | Eucalyptol |
| 41 | modd08_odour4 | Ethyl Tiglate |
| 42 | modd08_odour5 | Undecanal |
| 43 | modd08_odour6 | R-Citronellic Acid |
| 44 | modd15_odour1 | 4-Methyloctanoic Acid |
| 45 | modd15_odour2 | Ethyl Acetate |
| 46 | modd15_odour3 | 4-Methylacetophenone |

**Supp Table 1: List of monomolecular odours in the stimulus panel for each sample.**

| sample | animal ID | genotype info | gender | age | slice info |
|---|---|---|---|---|---|
| **C432** | ASAM3.2g | M72-ChR2-YFP, MOR174/9 GFP, Tbet-GCaMP6f | female | 11w4d | left bulb, 1st dorsal slice, 0.6 mm-thick |
| **Y391** | ASBC21.2a | MOR174/9 GFP, Tbet-GCaMP6f | male | 9w6d | left bulb, 1st dorsal slice, 0.5 mm-thick |
| **Y489** | ASBC21.2a | MOR174/9 GFP, Tbet-GCaMP6f | male | 11w4d | left bulb, 1st dorsal slice, 0.5 mm-thick |

**Supp Table 2: Mouse genotype, age and gender details of each sample.**

| sample ID | C432 | Y391 | Y489 |
|---|---|---|---|
| beam energy | 17 keV | 33 keV | 33 keV |
| temperature | cryo | cryo | room temperature |
| distances | dist 1, 2, 3, 4 | dist 2, 3, 4 | dist 1, 2, 3, 4 |
| step size | 1600 projections | 2000 projections | 2000 projections |
| exposure time | 300 ms | 300 ms | 150 ms |
| rotation angle | 180 degrees | 180 degrees | 180 degrees |
| acquisition mode | step with random jitter | step with random jitter | continuous |
| denoising | F | F | T |
| transmission | 14% | 52% | 60% |
| no. tiles - xy*z | 14 x 2 = 28 tiles | 16 x 2 = 32 tiles | 18 x 2 = 36 tiles |
| time per tile | 3h 15min | 2h 30min | 55min * 2 instances |
| total imaging time | 91 h | 80 h | 66 h |
| volume per tile | 0.026 $mm^3$ | 0.026 $mm^3$ | 0.026 $mm^3$ |
| total imaged volume | 0.728 $mm^3$ | 0.832 $mm^3$ | 0.936 $mm^3$ |
| high resolution volume per tile | 0.006 $mm^3$ | 0.006 $mm^3$ | 0.006 $mm^3$ |
| total imaged high resolution volume | 0.168 $mm^3$ | 0.192 $mm^3$ | 0.216 $mm^3$ |
| tissue containing volume | 0.21 $mm^3$ | 0.27 $mm^3$ | 0.30 $mm^3$ |
| sample geometry | parallelogrammic prism | parallelogrammic prism | cylinder |
| sample size | 965 x 1230 µm | 1320 x 1410 µm | diameter 1010 µm |

**Supp Table 3: XNH imaging parameters and data sizes.**

| counts | description | C432 | Y391 | Y489 |
|---|---|---|---|---|
| **curated 2P ROIs** | 2P cellular ROIs after removing duplicates and splits | 156 | 195 | 414 |
| **mapped cells** | curated 2P ROIs that were mapped to cells in XNH | 137 | 79 | 360 |
| **daughter cells** | mapped cells that were traced to parent glomeruli | 72 | 51 | 289 |
| **cell type** | | MC | TC | TC, MC |

**Supp Table 4: Yield of correlative experiments.**
